# Niche exclusion of a lung pathogen in mice with designed probiotic communities

**DOI:** 10.1101/2024.02.02.578711

**Authors:** Kelsey E Hern, Ashlee M Phillips, Catherine M Mageeney, Kelly P Williams, Anupama Sinha, Hans K Carlson, Kunal Poorey, Nicole M Collette, Steven S Branda, Adam P Arkin

## Abstract

For years, the airway microbiota have been theorized to be gatekeepers of respiratory health, as pathogens entering the airway make contact with resident microbes prior to or coincident with their interaction with host cells. Thus, modification of the native airway community may serve as a means of altering the local environment in favor of health. In this work, we hypothesize that synthetic bacterial communities introduced into the airway can serve as prophylactic countermeasures against infection by *Burkholderia thailandensis* in mice. We demonstrate that understanding of antagonistic interactions between a pathogen and airway microbiota *in vitro* can guide identification of probiotics with protective capabilities *in vivo*. Specifically, we show that niche overlap between the probiotic and pathogen is indicative of probiotic performance *in vivo*. This work serves as a foundation for the rational design of probiotic communities for protection against and treatment of respiratory infections.

## Introduction

Every mammalian body compartment that opens to the outside world has microbial life associated with it^1^. Until recently, the healthy lung was thought to be a sterile environment; however, recent work has demonstrated that the lower respiratory tract is colonized by a complex and roughly stable microbiome that impacts the health of the host^2–4^. For years, the airway microbiota have been theorized to be gatekeepers of respiratory health, as pathogens entering the airway make contact with its microbiota prior to or coincident with their interactions with host cells^3^. While the latter interactions have been studied in detail, little is known about pathogen-microbiome interactions in the airway.

It has been hypothesized that pathogen establishment in the airway could be halted by its microbiota through either indirect or direct competition with the pathogen^3,5,6^. During indirect competition, airway microbiota would modulate the host immune response to increase surveillance, promoting rapid recognition and clearance of foreign microbes including pathogenic organisms. During direct competition, the airway microbiota would competitively exclude a pathogen through production of inhibitory specialized metabolites, and/or by occupying a nutritional niche that would otherwise be used by the pathogen. Enhancement of these defense mechanisms might be achieved through addition of health-promoting microbes (probiotics) directly to the lower airway, an approach that has been demonstrated to prevent pneumococcal and *Pseudomonas aeruginosa* infection previously^7,8^.

While these studies have shed light on the utility of lower airway probiotics for preventing two specific infections, greater understanding of the principles that dictate probiotic efficacy would allow rapid identification of probiotics with therapeutic potential against many pathogenic targets. To enable development of these therapies in the future, pipelines for probiotic nomination need to be made which can rapidly and cheaply identify effective therapies. Further, systematic analysis of the mechanism of action of these probiotics would allow fine-tuned control of their activity allowing for more reliable therapeutic outcomes in the clinic.

Here, we seek to fill this gap in knowledge by systematically characterizing factors that make efficacious lower airway probiotics that can competitively exclude a model pathogen, *Burkholderia thailandensis* (*Bt*), from the lung environment. Specifically, we hypothesize that negative microbial interactions between a probiotic and pathogen better protect the host against infection. Our systematic study of the microbial ecology of these organisms enables development of a pipeline for nomination of effective single and multi-organism probiotics that, to this point, remain underdeveloped. We show that using our pipeline, we can accurately identify pairwise combinations of organisms with enhanced activity against *Bt*. Finally, we test the ability of these prophylactically administered probiotics to promote survival during a challenge with *Bt in vivo*.

## Results

### Isolation and characterization of candidate probiotics

To develop a system for testing the efficacy of lower airway probiotics in preventing infection, it was first necessary to identify a suitable pathogen to investigate. *Burkholderia thailandensis* (*Bt*) is a pathogen of mice that is genomically similar to several human pathogens including *Burkholderia pseudomallei, Burkholderia mallei* and organisms of the *Burkholderia cepacia complex,* the latter of which poses a high risk of mortality for cystic fibrosis patients^9–13^. Thus, *Burkholderia thailandensis* is an appropriate subject of study for our purposes as it is safe, infectious in mice and a good model of several pathogens with clinical relevance.

To acquire candidate probiotics for use in the lower airway, we sought organisms which would be able to survive in the lower airway for an extended period of time and be capable of tolerating its low-nutrient environment^14^. We hypothesized that organisms directly isolated from the airway would fulfill this criterion, as they are already well-adapted to the environment and may be more immunologically tolerated by the host. To isolate candidate probiotics, the trachea and lungs were collected from C57Bl/6j mice, homogenized, and plated onto five different solid growth media, either directly or after an initial period of culture in blood bottles. Tissues that should be sterile (liver and spleen) were collected from the same mice and processed in parallel, in order to recover any contaminants [“background” (B) strains] inadvertently introduced through our bacterial isolation procedures. Bacterial colonies were repeatedly streaked on LB agar to recover clonal isolates for whole-genome sequencing. In total, 28 isolates recovered from the lower airway tissues were further characterized for use as airway probiotics.

Prior work on the lung microbiome has led to the widely held hypothesis that many of its constituents originate from the oral and/or nasal microbiome and are unable to actively replicate and persist in the lower airway^15,16^. We therefore sought to identify airway isolates that are capable of active growth in the lower respiratory tract, with the supposition that they should be better equipped to re-colonize the airway and compete against an invading pathogen. To do this, the airway isolates were cultured in lung simulating medium (LSM), which is based on the nutrient composition of sputum samples from cystic fibrosis patients, and therefore is thought to approximate the nutritional environment of the lower respiratory tract^17–20^. We found that 10 of 28 airway isolates tested were capable of robust growth (>1 doubling within 48 hours) in LSM (Figure S1A). These 10 isolates, and model pathogen *Bt*, are arrayed on a phylogenomic tree (Figure 1A), and are henceforth referred to as candidate probiotics (CPs).

**Figure 1.**
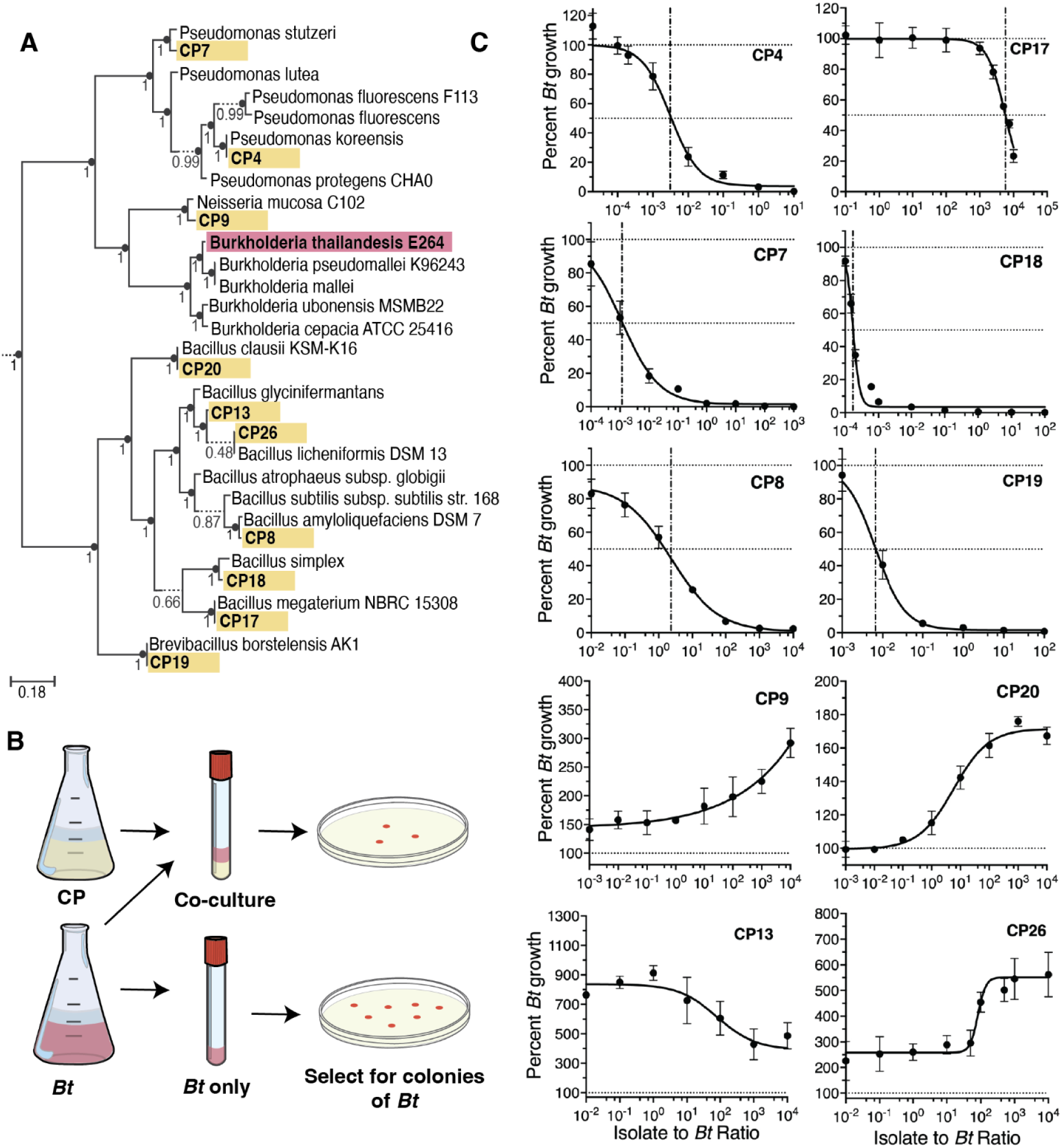
Characterization of CP activity against *B. thailandensis* in vitro. (A) Phylogenomic tree of CPs (yellow) and model pathogen *8. thailandensis* (Bt) (red), interspersed amongst publicly available genomes (no color). (B) Schematic of competition assay used to test in vitro inhibitory capabilities. Percent growth of *Bt* is calculated as the CFU in co-culture at 24 hours divided by the CFU in the *Bt* monoculture at 24 hours multiplied by 100. Co-culture inhibition was tested for multiple starting densities of CP (see Figure 1C x-axis). Starting density of *Bf* and CP at a 1:1 ratio are 3.33×104 CFU/ml per organism. *Bt* monoculture is inoculated at 3.33×10^4^ CFU/ml. (C) Dose-response curves for CPs against Bt. Relative IC50 is denoted with a vertical dashed-dotted line, and 100% and 50% growth are denoted with horizontal dotted lines. Percent growth at each density is represented as mean± SEM.

The phylogenetic assignments of the CPs were compared to those of airway microbiome constituents previously identified through 16S rRNA profiling of mouse airway tissue^21–25^. This analysis revealed that the taxonomic groups to which the CPs belong are well represented in the mouse airway microbiome (Tables S1-3). For example, we found that all 9 of the species to which the CPs belong were detected in at least 1 of 4 previous airway microbiome profiling efforts, with 7 of the 9 species detected in more than 1 profiling effort. In contrast, only 1 of the 9 species to which the B strains (contaminants recovered from liver and spleen) belong was detected, in only 1 of the 4 profiling efforts. Similarly, 3 of the 4 genera to which CPs belong were detected in all 4 profiling efforts (whereas for B strains, only 3 of 9 genera were detected in all 4 efforts); and all 4 of the families to which the CPs belong were detected in all 4 profiling efforts (whereas for B strains, only 6 of 8 families were detected in all 4 efforts). These results indicate that the CPs belong to taxonomic groups that are typically well represented in mouse airway microbiome, consistent with the idea that they are airway microbiome constituents that have been recovered into culture.

Next, we aimed to identify CPs capable of inhibiting *Bt* during co-culture in LSM, with the hypothesis that they might also inhibit *Bt* in the lower respiratory tract and therefore serve as efficacious probiotics. Expression of antagonistic phenotypes is dependent on several factors including the cell density relative to the available resources^26,27^. Accordingly, we decided to assay interbacterial interactions in LSM across varying cell densities, with the hypothesis that this would enable us to capture information about CP potency. Each CP was co-cultured with *Bt* in LSM at different relative ratios (Figure 1B). We found that the CPs showed a range of activities, including negative interactions (Figure 1C top 6 plots) and positive interactions (Figure 1C lower 4 plots) with *Bt*; the potency of each CP, as indicated by IC50 value, is shown in Table 1. Additionally, co-culture with *Bt* impacted growth of some CPs but not others, potentially affecting their interaction dynamics (Figure S1B).

**Table 1.**
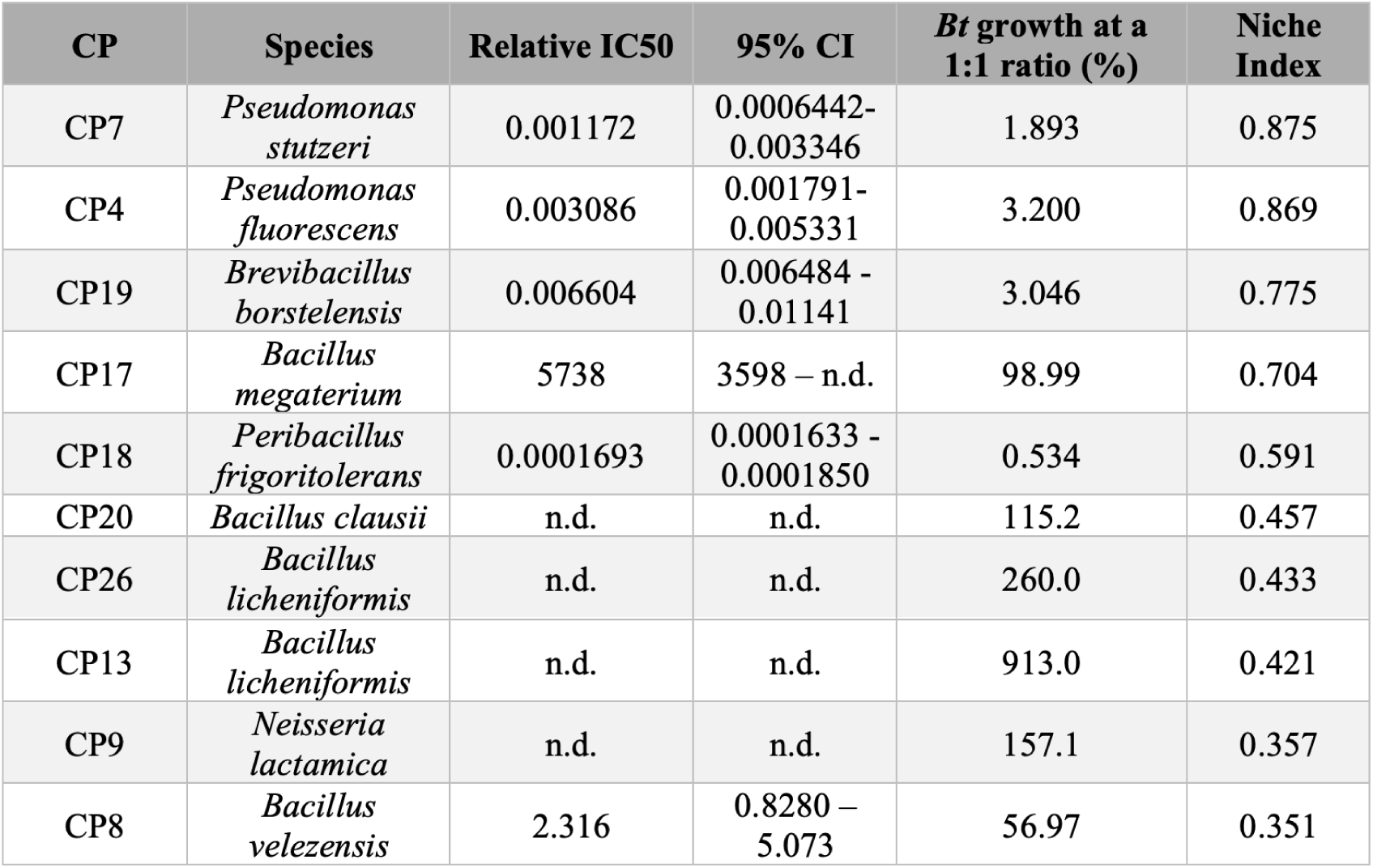
Summary of pathogen inhibition by CPs. Relative IC50 values and 95% confidence intervals (95% CI) are shown for each inhibitory CP. Not determined (n.d.) indicates that a value could not be measured. CPs are listed in order of decreasing NI values.

### Production of a specialized secondary metabolite plays a role in the antagonistic activities of CP8

Next, we sought to better understand the mechanism(s) of the observed negative interactions. First, we hypothesized that those CPs that strongly inhibit *Bt* may do so by simply growing at a faster rate than the pathogen thus dominating the environment. However, we found no correlation between doubling time of the CPs in LSM and their *Bt* antagonism (r = −0.21, P = 0.155, 95% CI [−0.47 to 0.088], N = 49). It has been shown previously that bacteriophages can modulate interbacterial competition, and some that can target *Bt* have been identified^28–31^; therefore, we next hypothesized that some of the negative interactions between CPs and *Bt* might be mediated through production of *Bt*-targeting phages. To test this, we attempted to recover phages from each CP using standard methods^32,33^, and tested the resulting product against *Bt* in plaque assays (Figure S2). We found that none of the CPs appeared to produce a phage that could effectively lyse *Bt*.

Given this information, we next hypothesized that these negative interactions could be direct, through chemical attack (ie. production of inhibitory metabolites); or indirect, through occupying a shared metabolic niche with the pathogen. First, we investigated the presence of inhibitory metabolites by growing *Bt* in sterilized co-culture supernatant from itself and a CP. We predicted that if *Bt*-inhibiting secondary metabolites were produced by a given CP during co-culture in LSM, then the conditioned medium recovered from the co-culture would inhibit growth of *Bt*. To test this, each negatively-interacting CP was co-cultured with *Bt* for 72 hours, filter sterilized, and combined with double-concentrated LSM to serve as medium for *Bt* growth curves (Figure 2A). We found that the co-culture supernatant from only one of the airway isolates (CP8) significantly reduced growth of *Bt* compared to supernatant from a *Bt* monoculture (negative control) (Figure 2B). This indicated that CP8 likely produced a specialized secondary metabolite and was engaging in interference competition. To confirm our findings, we performed agar diffusion assays on all inhibitory CPs and found that, aside from CP8, none produced a zone of inhibition on a lawn of *Bt* (data not shown).

**Figure 2.**
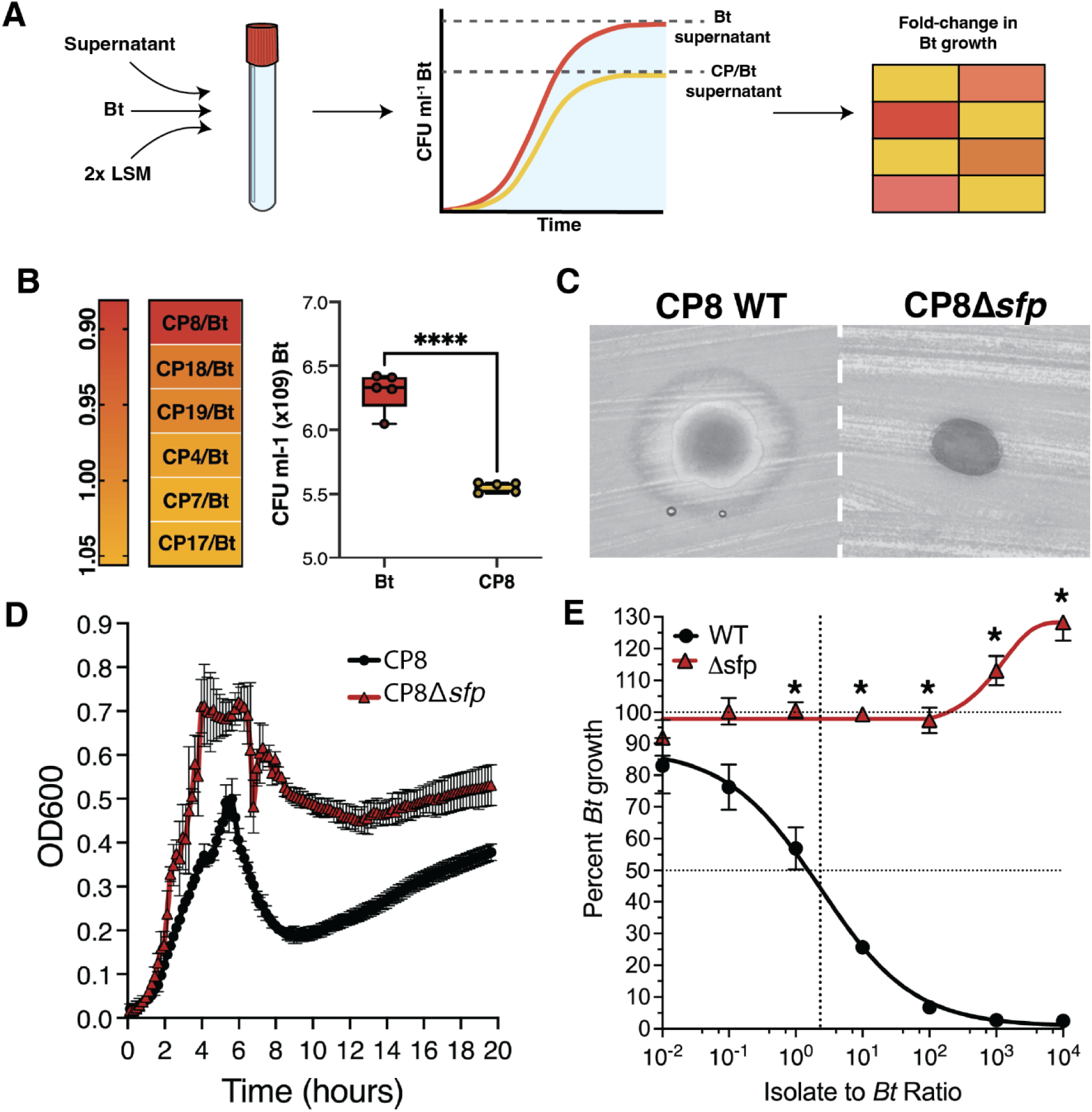
Production of a specialized metabolite plays a role in the antagonistic activities of CPS. (A) Schematic representation of supernatant inhibition experiment workflow. Supernatant from a *Bf* and CP co-culture or from *Bf* monoculture (supernatant) is mixed 1:1 with 2x LSM. This mixture is used as the growth medium for subsequent *Bf* growth curves. Fold-change in *Bt* growth is calculated by dividing the maximum growth of *Bt* in co-culture supernatant by the maximum growth of *Bt* grown in supernatant from itself. (B) Left: Heatmap of fold-change in max CFU of *Bt* grown in supernatant from each inhibitory CP and *Bt* co-culture relative to *Bt* grown in supernatant from itself. Co-culture supernatant was collected at 72 hours. A fold-change of 1 indicates no difference in growth in co-culture supernatant compared to *Bt* growth in supernatant from itself. A fold-change less than 1 indicates a reduction in pathogen growth. Right: Boxplot of growth inhibition by CPS supernatant. Data are represented as mean± SD. (C) Agar diffusion assays testing the CPS wildtype strain (CPS WT) and sfp mutant strain (CP8Δsfp) for inhibition of Bt after 48 hours. (D) Growth curves of CP8 WT (black) and CP8b.sfp (red) (E) Dose-response curve of CP8 WT (black) and CP8b.sfp (red) against Bt after 24 hours of co-culture. Percent growth at each density is represented as mean ± SEM.

To understand the degree to which CP8’s specialized metabolite is responsible for mediating the competitive phenotype observed in Figure 1C, we aimed to create a knockout of this molecule in CP8. Previously, we used antiSMASH^34^ to identify biosynthetic gene clusters in the CP8 genome sequence, and found that CP8 likely produces 19 specialized metabolites^35^. A review of the literature revealed that biosynthesis of many of these specialized metabolites is dependent on Sfp, a phosphopantetheinyl transferase which catalyzes their conversion to the active holo-form^36–39^. Therefore, we hypothesized that knocking out the *sfp* gene in CP8 would yield a mutant with an inability to produce a specialized metabolite with inhibitory capabilities against *Bt*. A clean deletion of the *sfp* gene was made in CP8, and the resulting mutant (CP8Δ*sfp*) was found to produce a smaller zone of inhibition in an agar diffusion assay, indicating that interference competition was reduced (Figure 2C). Additionally, growth curves of wildtype CP8 and the CP8Δ*sfp* mutant were performed, which showed no growth defect in the mutant strain (Figure 2D), indicating that its reduction in interference competition was not due to a growth defect. Finally, a competition assay was performed using the CP8Δ*sfp* mutant. The mutant showed reduced inhibitory activity compared to wildtype CP8 and, in contrast, demonstrated a positive interaction with *Bt* when interference competition was disrupted (Figure 2E).

### Metabolic niche overlap is indicative pathogen and CP antagonism

While production of an inhibitory metabolite appeared to be the primary mode of inhibition for CP8, 5 other CPs showed a negative interaction with *Bt* but did not appear to produce specialized metabolites. Previous work has shown that competition for a shared metabolic niche is an important mediator of interbacterial interactions in bacterial communities^40–44^. Therefore, we hypothesized that inhibition of *Bt* by these 5 CPs may be due to metabolic niche overlap. Specifically, we postulated that passive consumption of a limiting resource by the CP could reduce the supply to *Bt,* thus inhibiting its growth.

The Redfield Ratio describes the stoichiometric relationship between carbon, nitrogen and phosphorus which supports life on earth and is canonically 106:16:1^45^. Recent work has shown that this ratio is variable across body sites, with areas like the mouth and gut being more carbon rich, and the airways being relatively carbon poor^46^. Similar to the upper airway, LSM is carbon limiting, with a C:N:P ratio of approximately 88:14:1. This suggests that carbon may be the limiting nutrient in the lower airway and thus, may be the limiting resource in the airway niche. We predicted that if carbon was limiting, then supplementation of LSM with additional carbon would reduce the antagonism between the CP and *Bt*. To test this, LSM was supplemented with 43mM glucose to increase the C:N:P ratio to 101:14:1. We observed a leftward shift of the dose-response curve for CP7 in the additional carbon condition (Figure 3A), and a significant reduction in inhibition at a 1:100 ratio (Figure 3B). A similar result was observed when the same experiment was conducted with CP19 (Figure S3). These results suggest that with increased carbon abundance there is a reduction in antagonism, consistent with the idea that, under the carbon limiting conditions of the airway, carbon is the primary nutrient for which CPs and *Bt* are competing.

**Figure 3.**
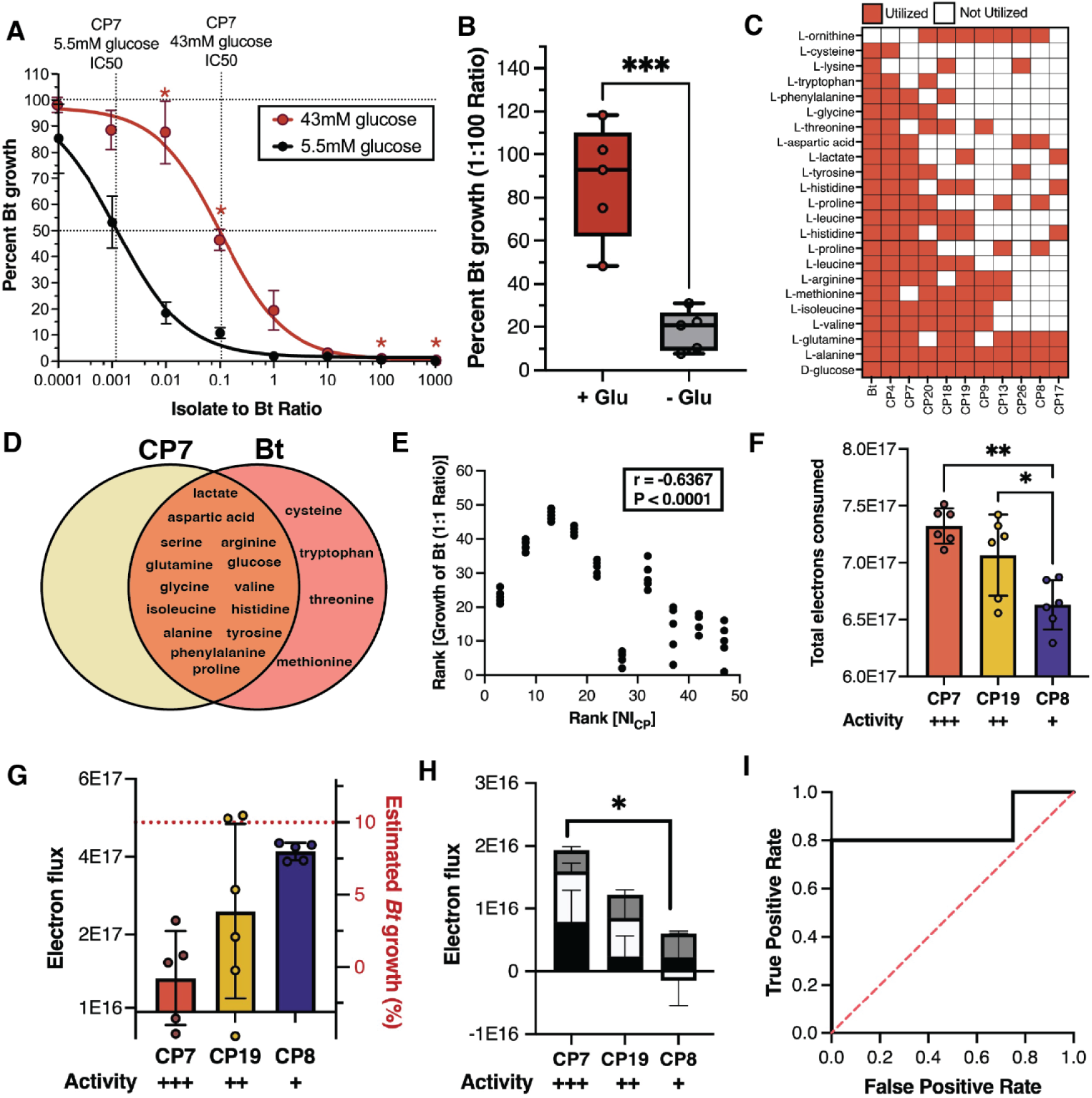
Metabolic niche overlap is indicative of pathogen and CP antagonism. (A) Dose-response curve for CP7 grown in unsupplemented LSM (black) or with additional glucose (red). Percent growth at each density is represented as mean ± SEM. (B) Activity of CP7 against Bt at a 1:100 ratio with or without glucose supplementation (red and gray, respectively). Data are summarized as mean ± SD. ***P<0.001. (C) Carbon utilization heatmap for CPs and Bt. (D) Venn diagram of carbon sources utilized by CP7 and Bt. (E) Plot of rank percent *Bt* growth at a 1:1 ratio after 24 hours vs rank of Niche Index for each CP. Correlation is determined by Spearman rank correlation (r = −0.6367, P < 0.0001, 95% Cl [−0.7891 −0.4261], N = 49). (F) Total carbon consumed by CPs grown in combination with Bt at 36 hours (mean± SD). Activity indicates relative amount of Bt inhibition at a 1:1 ratio from most(+++) to least(+) inhibitory. *P=0.05; **P<0.01. (G) Total electron flux at 24 hours. Red dashed line indicates the estimated abundance of *Bt* in each co-culture. (H) Summed electron flux of lactate (black), proline (white), and aspartic acid (gray) at 24 hours in different CP co-culture conditions (mean± SD). *P<0.05.(I) Receiver operator characteristic curve showing the ability of growth on lactate to distinguish inhibitory CPs from non-inhibitory CPs. An AUROC of 1 indicates perfect distinction between inhibitors and non-inhibitors; an AUROC of 0.5 (dashed red line) indicates no distinction between inhibitors and non-inhibitors.

Further, we sought to explore the extent to which metabolic overlap with *Bt* for carbon sources could predict inhibitory capabilities of each CP. To do this, we assayed the growth of each CP and *Bt* in each individual carbon source from LSM to determine which carbon sources could be utilized by each organism (Figure 3C). *Bt* showed a wide range of metabolic potential and was able to grow on every carbon source presented except for ornithine. This is perhaps unsurprising as it has been shown previously that pathogens acquire new metabolisms in order to survive in their host^47^. Conversely, the CPs displayed diverse metabolic activity, with CP4 and CP7 appearing most similar to *Bt* (Figure 3C).

Each carbon source in LSM is used both for production of biomass as well as the primary electron donor to generate energy via aerobic respiration. In the niche exclusion hypothesis, a competitive advantage can be obtained for an organism that is able to consume those resources that would otherwise fuel aerobic respiration of a competitor. We therefore hypothesized that those CPs that are capable of sharing a greater proportion of carbon-derived electrons with *Bt* would have more potent antagonism of the pathogen by covering a greater percentage of the energetic niche (Figure 3D). On this assumption, we built a simple index of niche overlap which weights each carbon source in LSM utilized by a given CP by the theoretical electron contribution of that given source, with the belief that such a metric could identify probiotics capable of achieving maximal niche coverage. We hypothesized that weighting each carbon source by its theoretical electron contribution would improve the accuracy of the niche index, as those sources which contribute more electrons represent a greater proportion of the energetic niche and therefore have a potentially greater impact on antagonistic phenotypes. While many excellent models exist for metricizing niche overlap^48–50^ and greatly inspired our model design, we sought to create a model which would describe the metabolic niche of the respiratory tract as a function of the energy available to the microorganisms within it.

Niche Index (NI) is calculated as follows:

First, the number of electron equivalents generated by each carbon source is calculated assuming the complete oxidation of these sources to carbon dioxide under standard conditions. The electron equivalents are then multiplied by the total number of molecules of the carbon source in the media. This yields the total electron equivalents for each carbon source per liter of media. These values are held in vector 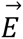.

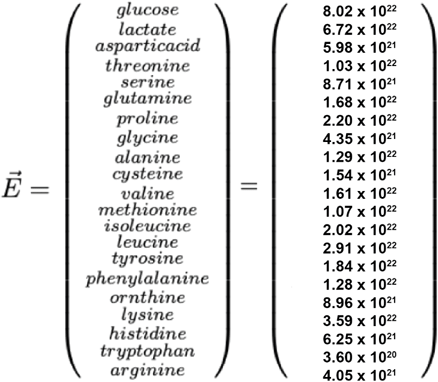

The weighted value for each carbon source is:

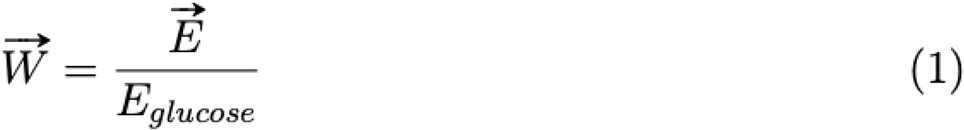

For all CPs and *Bt*, a carbon utilization vector is generated. If a carbon source is consumed, then i = 1, if it is not able to be consumed, then i = 0.

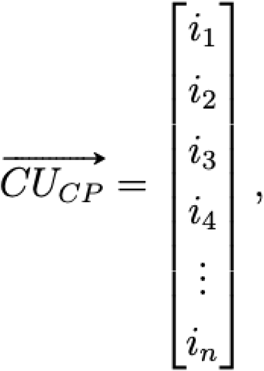

To calculate the Niche Index of each CP with *Bt*, the Hadamard product of the carbon utilizations vectors for each CP and *Bt* are multiplied by the weighting vector. The resulting value represents the electron equivalents utilized by both the CP and *Bt*. This value is divided by the electron equivalents utilized by *Bt* alone. Thus, the equation for Niche Index is: :

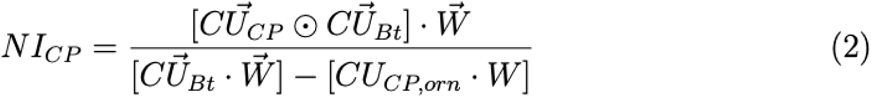

NI is higher when a given CP shares a greater number of carbon-derived electrons with *Bt.* To evaluate the association between the co-culture interaction with *Bt* and the NI of each CP, a non-parametric spearman rank-order correlation was performed with the null hypothesis that there is no monotonic association between the two variables. We found that NI is negatively associated with the co-culture growth of *Bt* with a given CP at a 1:1 ratio (r = −0.64, P < 0.0001, 95% CI [−0.79 −0.43], N = 49) (Table 1 and Figure 3E). Since CP8’s potency is primarily due its secondary metabolite production (Figure 2E), percent growth of the pathogen in co-culture with the CP8Δ*sfp* mutant was used in this analysis to understand how NI relates to CP8’s interaction with *Bt* (Figure 3E). While here we have used Niche Index to calculate niche overlap between CPs and *Bt* in LSM, this model is generalizable and can be used to calculate the niche overlap between organisms in any media with a known molar carbon composition.

To confirm our hypothesis that those organisms that consume a greater number of carbon-derived electrons with *Bt* are more antagonistic, we performed exometabolomics on supernatants from 3 CPs. These 3 CPs represent a range of inhibition strengths and niche indexes, with CP7 being the most potent (IC50 = 0.001172 NI = 0.875), CP19 displaying moderate potency (IC50 = 0.006604 NI = 0.775) and CP8 being the least potent (IC50 = 2.316 NI = 0.351). Each of these CPs was combined with *Bt* at their respective IC90 and co-incubated for 36 hours. Supernatants from these CPs at 36 hours show that more potent CPs consume a statistically greater amount of carbon from the media than less potent CPs (Figure 3F). At 24 hours where the degree of inhibition of *Bt* by each CP is the same, more potent probiotics consume fewer carbon-derived electrons than less potent competitors (Figure 3G). This led us to hypothesize that certain carbon sources were being prioritized, allowing more antagonistic CPs to have the same net effect on *Bt* while consuming fewer electrons. Further, we speculated that if high priority carbon sources were to be identified, perhaps the consumption of a smaller set of these high priority carbon sources could predict activity with similar accuracy to the Niche Index.

From the exometabolomics data, consumption of three carbon sources, lactate, proline and aspartic acid are prioritized more at 24 hours by the most potent candidate probiotic, CP7, than the least potent, CP8 (Figure 3H). Furthermore, we found that a CP’s ability to generate biomass on lactate (the second most abundant carbon source in LSM) was a good predictor of its ability to inhibit the pathogen (Figure 3I), suggesting that the ability to consume lactate in co-culture may be important for pathogen control. Overall, these results indicate that consumption of specific resources can have an especially strong influence on niche exclusion, with consumption of a specific carbon source (lactate) appearing to play a key role in niche exclusion of *Bt* by the most potent of the CPs (CP7). While these results are promising building blocks for the development of a metric of activity based upon a smaller feature set, a larger screen of candidate probiotics may be necessary to understand the generalizability of these features. Further, future efforts to develop this model should take into account a CP’s uptake rate for a given resource as this may influence antagonism and reveal features of the metabolic niche that have disproportionate significance to interbacterial interactions.

### Niche overlap aids in the identification of efficacious multi-organism probiotics

While our work showed that 6 of the CPs were potent inhibitors of *Bt* growth in co-culture, 4 were poor inhibitors or actually promoted *Bt* growth (Figure 1). Having found that NI is well correlated with CP activity *in vitro*, we hypothesized that this metric could be altered to identify pairwise combinations of poor-performing CPs with enhanced antagonism. To do this, we created a new metric, the Niche Index Fraction (NIF), which is calculated by generating the NI between two CPs and dividing this value by the NI of the two CPs and *Bt* (Figure 4A). We hypothesized that combinations with lower NIF values would have optimal coverage of the *Bt* niche space while minimizing niche overlap between two CPs. Thus, low NIF values might indicate combinations with enhanced inhibition of the pathogen compared to each CP alone. NIF is calculated as follows:

**Figure 4.**
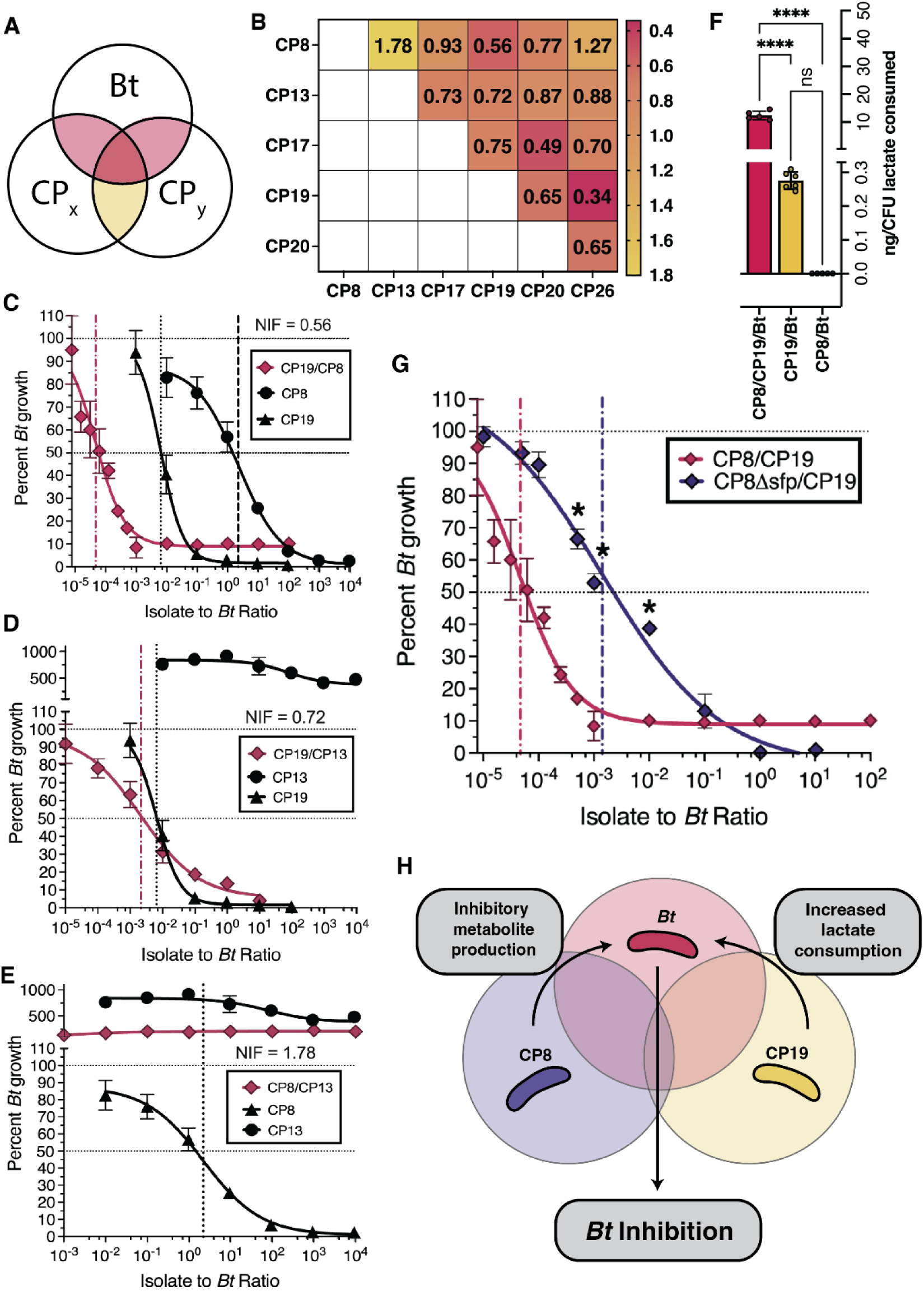
Niche overlap aids in identification of efficacious CP combinations. (A) Venn diagram representation of the Niche Index Fraction (NIF) calculation. Yellow represents the fraction numerator and red the denominator. (B) Heatmap of NIF values for each colonizing CP. (C) Dose-response curve for the CP8/CP19 combination (mean ± SEM). (D) Dose-response curve for the CP19/CP13 combination (mean ± SEM). (E) Dose-response curve for the CP13/CPB combination (mean± SEM). (F) Lactate consumed (as determined via GC-TOF) for 3 conditions, normalized to CFU of each CP inoculated (mean ± SD). ****P<0.0001. (G) Dose-response curve for the CP8/CP19 (red) and CP8sfp/CP19Δsfp (black) combinations. (H) Schematic of the proposed mechanism for *Bt* inhibition by the CP8/CP19 combination. CP8 produces a specialized metabolite that inhibits growth of the *Bt.* Additionally, increased lactate consumption by CP19 in the presence of CP8 further reduces *Bt* growth. Overall, minimal niche overlap between the CPs, and high overlap with *Bt,* allows for further inhibition of *Bt* with minimal inter-CP antagonism.

First, the overlap between two CPs is generated and is weighted by the relative electron contribution of each carbon source

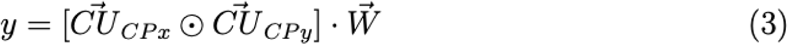

Total carbon-derived electron utilization is calculated for both CPs

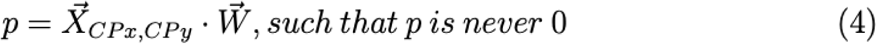

For which

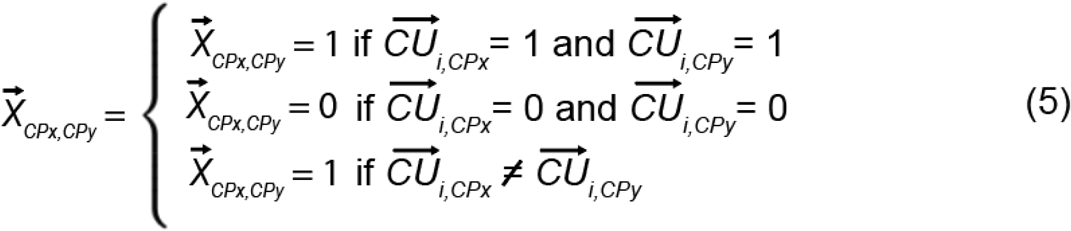

Thus the niche overlap between two CPs is

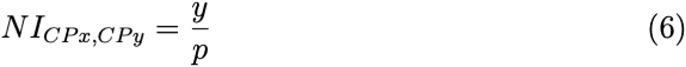

The niche overlap between two CPs and *Bt* is thus

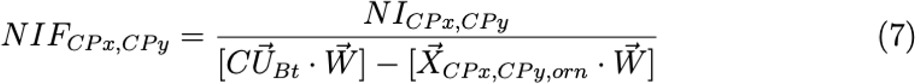

We calculated NIF for every pairwise combination of the 7 least antagonistic CPs from Figure 1. This generated a matrix of CP combinations and their respective NIF values (Figure 4B). To test this metric’s capability to identify inhibitory combinations of CPs, we chose to evaluate pairwise combinations with the three lowest NIF values, a moderate value and a high value using competition assays.

In these experiments, two CPs were combined at a 1:1 ratio such that the total CP concentration at a given ratio was the same as their individual CP counterparts. This enabled us to directly compare activity of an individual CP and the activity of a combination of two CPs at a given ratio as the total cell density was the same in each. From competition assays, we determined that NIF was able to predict combinations with enhanced and reduced *Bt* antagonism as reflected by their relative IC50 values (Figure 4C,D,E, Figure S4 and Table S4).

For one combination, CP8 and CP19, we sought to understand the mechanism by which antagonism was being enhanced. First, to understand the effect of the interaction on all members of the co-culture, we performed growth curves of each strain (CP8, CP19 and *Bt*) in co-culture and compared their individual abundances when grown together to their growth in monoculture using qPCR (Figure S5). Reduced growth in co-culture of all three organisms in comparison to monoculture (Figure S1) classifies this antagonistic interaction as true competition. To better understand how CP8 and CP19 may enhance each other’s antagonistic impact on *Bt*, we performed exometabolomics analysis on supernatant from this combination. At 24 hours, significantly more lactate was consumed per cell by the CP8/CP19/*Bt* co-culture than by the CP8/*Bt* co-culture or CP19/*Bt* co-culture (p<0.0001) (Figure 4F). Similarly to the single isolate studies, this result once again suggests that lactate utilization may be important for antagonism of *Bt* in an airway-simulating environment.

To further investigate how the combination of CP8 and CP19 more effectively inhibits the pathogen, we performed a competition assay using the CP8Δ*sfp* mutant in combination with CP19 to understand the degree to which secondary metabolite production was responsible for their combined effect (Figure 4G). When wildtype CP8 was replaced with the CP8Δ*sfp* mutant, there was a significant reduction in *Bt* inhibitory activity compared to the wildtype CP8/CP19 combination. Together, these findings suggest that both consumption of specific carbon sources and secondary metabolite production are important for activity of the CP8/CP19 combination and likely enable *Bt* inhibition activity (Figure 4H). While the concept of NIF suggests that coverage of the pathogen niche may be responsible for the observed activity in other combinations, further investigation needs to be done to understand these interactions in more detail.

### CPs provide protection against infection

Having observed the ability of the CPs to compete with *Bt* in an *in vitro* airway-simulating environment, we were encouraged to test their ability to confer protection against respiratory *Bt* infection *in vivo*. We hypothesized that CPs that can colonize at high density over an extended period of time would provide better protection against *Bt* infection. To test the airway colonization capabilities of the CPs we administered each to C57Bl/6j mice at 10^6^ CFU via oropharyngeal aspiration (OPA) and after a period of 7 days we assessed the lungs and trachea for CP load via CFU enumeration (Figure 5A). We found that several CPs were able to colonize the airway for the duration of the 7-day period (mean airway CP load of >10 CFU) (Figure 5B).

**Figure 5.**
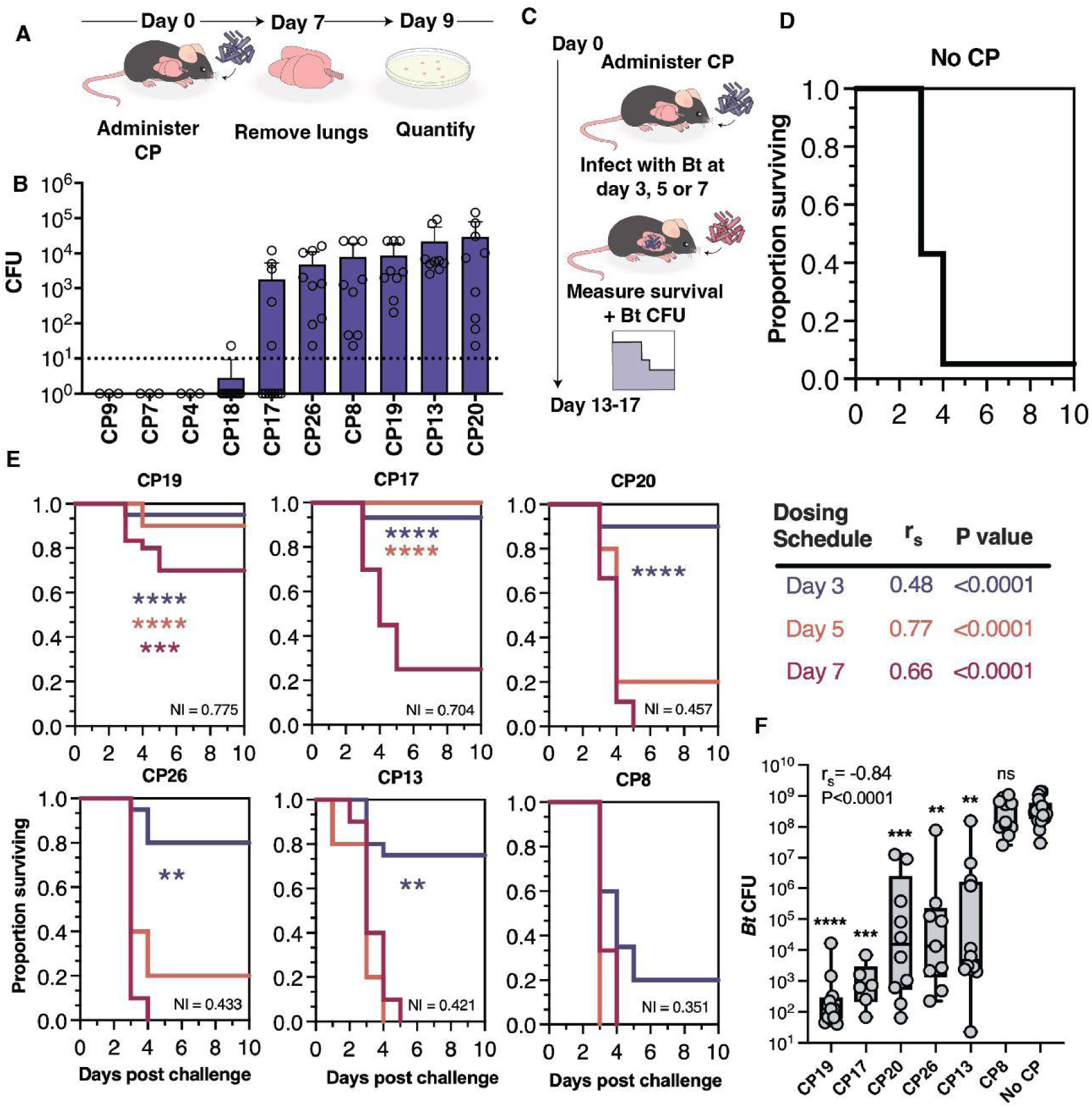
CPs colonize the mouse airway and protect against respiratory *Bt* infection. (A) Schematic of colonization testing for CPs. Each CP (10^6^ CFU) is administered to the lower airway via OPA. After 7 days, the airway tissues (lungs and trachea) are collected, homogenized, and plated in order to enumerate viable CPs (expressed as CFU per tissue homogenate) (B) Colonization results from airway tissues collected at 7 days following CP administration. The dotted line represents the minimum CFU at which the CP is considered able to colonize in a reliably detectable manner. Data are represented as the mean ± SD. (C) Schematic of the method for survival testing. Each CP (10^6^ CFU) is administered to the lower airway via OPA at 3, 5 or 7 days post-GP a normally lethal dose of of *Bt* (3×10^4^ - 5×10^5^ CFU) is administered to the lower airway via OPA. Survival is measured over the course of 10 days post-Bt. (D) Survival data from mice administered no CP (vehicle control) and challenged with *Bt* (E) Survival data from mice administered CPs at 3 days (blue), 5 days (orange), or 7 days (red) prior to *Bt* challenge. The Niche Index value for each CP is listed in the lower-right corner of its associated graph. Comparison of survival rate with versus without CP treatment was accomplished using the Mantel-Cox test ****P<0.0001, ***P<0.0002, **P<0.01. A summary of the relationship between days surviving and Niche Index value for each dosing schedule is also shown (F) Pathogen loads in airway tissues of mice prophylactically treated with CPs. Each CP or PBS (“no CP” negative control) is administered to the lower airway via OPA, 3 days later Bl (3×10^4^ - 5×10^5^ CFU) is similarly administered, and 3 days later (i.e., 3 days post-challenge, 6 days post-treatment) the airway tissues are collected for enumeration of Bl CFU. Mean *Bt* CFU counts are as follows for each group: CP19 (1.6×10^3^), CP17 (1.7×10^3^), CP20 (2.2×10^6^), CP26 (8.7×10^6^), CP13 (1.3×10^7^), CPS (3.9×10^8^), No CP (4.6×10^8^). Asterisks indicate significant differences in Bt CFU between treatment versus no treatment groups as determined using Dunn’s multiple comparisons test ****P<0.0001, ***P<0.0002, **P<0.01. The relationship between the Niche Index values of CPs and the Bl CFU detected in mice treated with the CPs is indicated in the upper left corner.

We screened both colonizing and non-colonizing CPs for their ability to confer protection in a mouse model of respiratory *Bt* infection. In these studies, 10^6^ CFU of each of 6 CPs were administered to the airway of C57Bl/6j mice via OPA. Following a period of 3, 5, or 7 days, *Bt* was administered to the airway via OPA and mortality was monitored for 10 days following *Bt* challenge (Figure 5C). Mice administered no CP prior to *Bt* challenge showed high mortality, only rarely surviving past day 4 (Figure 5D). In contrast, mice administered certain CPs prior to *Bt* challenge showed significantly reduced mortality (Figure 5E). Five of the six CPs were protective when administered 3 days prior to *Bt* challenge; CP17 was protective when administered at 3 or 5 days prior to *Bt* challenge; and CP19 was protective when administered at 3, 5, or 7 days prior to *Bt* challenge.

We observed that the most protective probiotics, CP19 and CP17, were predicted to have the highest niche overlap with *Bt* as indicated by their NI values (Table 1), showed an ability to inhibit *Bt* in competition assays (Figure 1C) and reduced *Bt* loads in mouse airway tissues during respiratory infection (Figure 5F). Furthermore, the CPs that robustly colonized the airway (Figure B) but did not inhibit *Bt* in competition assays (Figure 1C) (i.e., CP13, CP20, and CP26) showed no significant protection when administered at 5 or 7 days prior to *Bt* challenge (Figure 5E). A notable exception to this trend was CP8, which showed robust colonization as well as inhibition in competition assays, but showed no significant protection of mice challenged with *Bt*. This may be explained by its low niche overlap with the pathogen (Table 1) and/or its reduced colonization in *Bt*-infected mice (Figure S6). Taken together, these results suggest that niche overlap with the pathogen may be suggestive of a CPs protective capabilities. However, given the potential for certain CPs to inhibit *Bt* via the production of specialized metabolites (Figure 2), future work should be done to investigate the role of these unmodeled interactions *in vivo*.

We sought to estimate the strength of correlation between NI_CP_ and CP protection efficacy. Accordingly, we performed a non-parametric spearman rank correlation analysis with the null hypothesis that there existed no relationship between NI_CP_ and the number of days surviving when CP treatment was performed 3, 5 or 7 days in advance of pathogen challenge. We found a moderately strong relationship between NI_CP_ and survival when CPs are dosed 3 days prior to pathogen challenge (r = 0.48, P<0.0001, 95% CI [0.33 to 0.62], N = 115), a very strong relationship when dosed 5 days prior (r = 0.77, P<0.0001, 95% CI [0.60 to 0.88], N = 40), and a strong relationship when dosed 7 days prior (r = 0.66, P<0.0001, 95% CI [0.52 to 0.77], N = 89) (Figure 5E). Additional studies revealed that CP13, CP20, and CP26, which robustly colonized the airway (Figure 5B) but did not show inhibition in competition experiments (Figure 1C) and conferred a survival benefit only when administered 3 days prior to pathogen challenge (Figure 5E), provided similar protection even after being rendered non-viable (via UV and heat treatment) (Figure S7). In contrast, we found that the protective effects of CP17 and CP19 when administered 5 or 7 days prior to pathogen challenge largely depended on CP viability. Taken together, these results suggest that niche exclusion is the dominant mechanism of protection except when CPs are administered 3 days prior to pathogen challenge, in which case other mechanisms that are not captured in our model (potentially including immune priming) must play a role.

We further sought to determine whether the survival benefits conferred by CPs were associated with reduced pathogen loads in the airway, which would be consistent with the niche exclusion mechanism of protection. In these studies the CPs were administered to the airway 3 days prior to *Bt* challenge, and airway tissues collected at 3 days post-challenge for *Bt* CFU analysis. We found that prophylactic treatment with the CPs significantly reduced pathogen loads in the airway, with the notable exception of CP8 (Figure 5F). Moreover, we observed a very strong inverse relationship between NI_CP_ and pathogen load (r = −0.84, P<0.0001, 95% CI [−0.90 to −0.76], N = 77), where CPs with higher NI values more effectively reduced pathogen loads. While it appears that NI is a reasonable indicator of the protective effects of a CP *in vivo*, additional development of this model will need to be done to better capture the host environment, improve prediction, and determine the extent to which it can be used to nominate efficacious probiotics in the future.

From our *in vitro* studies, we found that lactate consumption by a CP was important for pathogen inhibition. To study the specific role of lactate in a scenario more similar to the *in vivo* context, CP19 (10^6^ CFU) or PBS (negative control) was introduced into airway tissue homogenates and after 24 hours of incubation the lactate levels were measured. We found that lactate levels were significantly lower in tissue homogenates inoculated with CP19 as compared to those receiving PBS only (Figure S8A). We also found that sterile filtrates recovered from CP19-conditioned tissue homogenates failed to support growth of *Bt*, in contrast to those recovered from PBS-treated tissue homogenates (Figure S8B). These results suggest a link between lactate depletion by CP19 and its ability to inhibit the pathogen in the context of airway tissues.

## Discussion

The human respiratory tract microbiome composition has been linked to both respiratory health and the occurrence of respiratory infections^51^. As a result of this connection, manipulation of the lower airway microbiome is an attractive strategy for anti-infective therapies. Airway microbiome modulation via supplementation with probiotics has previously shown success in preventing infections; however, these studies lack a more broad and systematic analysis which is necessary to learn generalizable principles for the design of efficacious airway probiotics^7,8^.

In this work, we developed a system that probes interactions between airway-derived candidate probiotics (CPs) and a model respiratory pathogen *Burkholderia thailandensis* (*Bt*). From this exploration, we find that consumption of shared resources explains the majority of antagonistic phenotypes between CPs and *Bt in vitro*. Specifically, consumption of shared carbon sources seems to play an important role in these interactions. As a result, a metric of niche overlap, Niche Index (NI), is correlated with CP inhibitory activity *in vitro* in most instances. Furthermore, we use these same principles to design combinations of CPs with enhanced antagonism against *Bt*. We explore the mechanism by which one combination, CP19 and CP8, exploits both metabolic niche exclusion and inhibitory metabolite production to its advantage in competing with *Bt*. Finally, we find that several CPs are able to re-colonize the mouse airway for at least 7 days, and that two CPs (CP19 and CP17) significantly reduce mortality in mice subjected to otherwise lethal respiratory *Bt* challenge. While it appears that there is a relationship between the NI and the protective capabilities of a CP, further preclinical experiments will need to be done to determine if NI alone can guide formulation of airway probiotics with clinical applicability. Further, consideration of potential effects from patient genetic background, environment, host immune response, and cross-feeding must also be explored.

The method described here is a step towards a more rapid, inexpensive, and effective means of identifying efficacious probiotics, all factors that have stalled probiotic development previously^52^. Selection of these probiotics based on their niche-exclusion capabilities, as demonstrated here, also has the potential to enable development of probiotics with better safety profiles. While probiotics with the ability to produce inhibitory small molecules can be identified easily, pathogens can rapidly evolve to evade this form of antagonism. In comparison, probiotics which occupy a niche may leave a pathogen with fewer strategies to evolve resistance, allowing long-term efficacy and more predictable effects on health in clinical applications.

Another great challenge for clinical development of airway probiotics is potency. In order to allow for adequate oxygen exchange in the lungs, alveoli need to remain unobstructed and, thus, colonization of the lung with a large dose of bacteria may result in undesirable outcomes. Therefore, probiotics which require extremely low doses to achieve a protective effect are desirable. Our methodology shows potential for repurposing of organisms that are typically poor performers for use in highly potent multi-organism formulations.

In this work, we found that one way that probiotics defend against a pathogen is through consumption of available carbon sources which would otherwise feed the pathogen. In particular, lactate consumption appears to be important for efficacy. Several studies have found that there is increased lactate in the lung during inflammatory exacerbations, which could feed pathogen growth^53–55^. Interestingly, previous studies have shown that consumption of lactate is important for fitness in the host and pathogenesis for several different bacterial pathogens^56–60^. Theoretically, if probiotics that preferentially consume lactate can be designed, they may be useful in controlling infections in hosts with inflammatory lung conditions, such as cystic fibrosis, idiopathic pulmonary fibrosis, and chronic obstructive pulmonary disease^54,55,61^.

Broadly, we hypothesize that airway probiotics may have functionality by stopping the positive feedback loop of infection. In this model, pathogen infiltration into the lung and growth results in inflammation and cellular injury, causing a leak of nutrient-rich fluids into the alveolar compartment, which further promotes pathogen growth (Figure 6 (left))^15^. Airway probiotics might stop this feedback loop by depleting nutrient abundance in the local lung environment, thereby limiting pathogen growth (Figure 6 (right))^15^. Given this model, probiotics which interfere with pathogen metabolism could be forward-designed to work optimally within these environments in order to maximize potency and efficacy.

**Figure 6.**
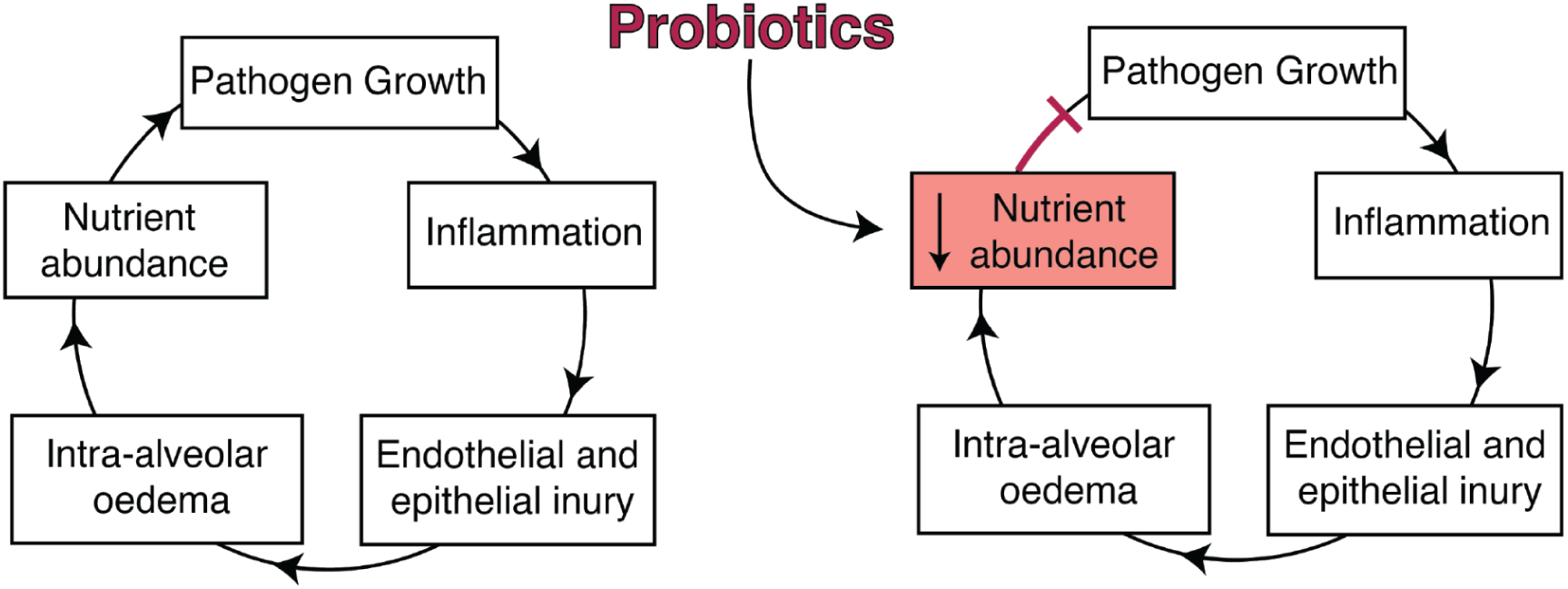
Proposed mechanism of CP protection. Left: During lower airway infection, bacterial growth elicits inflammation from the host, with concurrent endothelial and epithelial injury. Intra-alveolar oedema introduces additional nutrients from the blood into the alveoli and lung lumen, enabling further growth of the pathogen. This positive feedback loop results in uncontrolled growth of the pathogen, ultimately facilitating its instantiation. Right: Probiotics delivered directly to the lower airway may limit infection by reducing nutrient abundance after oedema, which prevents further growth of the pathogen and thereby breaks the infection-promoting feedback loop.

The native airway microbiome, and its interactions with host cells and incoming pathogens, is only partially understood, making development of airway probiotics a challenge. As our knowledge grows, health-promoting functionalities could be introduced into the lower respiratory tract, through supplementation with microbiome constituents in unaltered state (as demonstrated here) or with engineered properties, such as novel metabolisms. However, for utility in the clinic, numerous questions will need to be answered about the predictability, long-term safety and reliability of airway probiotics as alternatives or additions to antibiotic therapies. To improve this nomination pipeline further, future models should incorporate elements which have been shown to influence interbacterial antagonism previously such as the rate of consumption of shared resources and and the impact of sudden alterations in resource availability as might occur in acute lung injury. To ensure safety, evolution of the CP in the airway should be monitored over time, and the potential contribution of cross-feeding between CP and pathogen should be investigated. Additionally, future efforts will need to focus on reliable probiotic delivery modalities, formulation stability, and potential for variance in colonization, protection, and immunogenic responses in different hosts and environmental contexts. Work is currently underway to understand the generalizability of the findings here to models of other respiratory diseases, and to evaluate their safety and efficacy for therapeutic applications.

## Acknowledgements

We thank K. Sander (UCB) and K. Vyas (CMS) for helpful discussion; D. Sivanandan (UCB) for assistance with experiments; and M. Hirakawa (SNL) for critical review of the manuscript. We thank the West Coast Metabolomics Center for exometabolomics services. Sandia National Laboratories is a multimission laboratory managed and operated by National Technology & Engineering Solutions of Sandia, LLC, a wholly owned subsidiary of Honeywell International Inc., for the U.S. Department of Energy’s National Nuclear Security Administration under contract DE-NA0003525. This work was performed under the auspices of the U.S. Department of Energy by Lawrence Livermore National Laboratory under Contract DE-AC52-07NA27344.This work was performed under the auspices of the U.S. Department of Energy by Lawrence Livermore National Laboratory under Contract DE-AC52-07NA27344.

## Author Contributions

Conceptualization, K.E.H., S.S.B. and A.P.A.; Methodology, A.M.P., N.M.C., K.E.H. and H.K.K; Investigation, A.M.P, N.M.C., S.S.B., K.E.H. and H.K.K.; Formal analyses - C.M.M., K.P.W., K.P., S.S.B., A.S., and K.E.H.; Writing – Original Draft, K.E.H.; Writing - Review & Editing, all authors; Funding acquisition, S.S.B. and A.P.A; Supervision, S.S.B. and A.P.A.

## Methods

### Isolation and Culture of Bacteria

A *Burkholderia thailandensis* E264 strain that constitutively expresses GFP was provided by Daniel J. Hassett (University of Cincinnati College of Medicine; Cincinnati, OH)^62^. The candidate probiotic (CP) isolates were recovered from the lower respiratory tracts of healthy mice using a culturomics approach^63^. All animal work was conducted in accordance with protocols approved by the Lawrence Livermore National Laboratory (LLNL) Institutional Animal Care and Use Committee (IACUC, protocol 304) and Institutional Biosafety Committee (IBC, protocol 2021-010). LLNL is accredited by the Association for Assessment and Accreditation of Laboratory Animal Care (AAALAC) International, and is Public Health Service-Assured (PHS assurance A3184-01). Briefly, airway (trachea and lung) tissues were collected from 24 mice (C57Bl/6j, 10-20 weeks), homogenized in 500 uL of phosphate-buffered saline (PBS), and used to inoculate five different solid growth media [tryptic soy (TS); TS + 5% sheep’s blood; M9 minimal salts + 0.2% glucose; brain-heart infusion (BHI); and Luria broth (LB) (Teknova; Hollister, CA)] as well as three different BACTEC blood bottles [Plus Aerobic/F; Plus Anaerobic/F; and Lytic/10 Anaerobic/F (Becton Dickinson; Franklin Lakes, NJ)]. The inoculated solid growth media were incubated at 37C for seven days. The inoculated blood bottles were incubated with shaking (250 rpm) for 1-5 days, and 100 uL aliquots were periodically withdrawn for use in inoculating the five different solid growth media, which were then incubated at 37C for seven days. All colonies detected on the solid growth media were individually transferred to LB agar and, after further incubation at 37C for 1-3 days, streaked to single colonies on fresh LB agar. A representative single colony from each Petri plate was used to inoculate LB liquid and solid media, and these cultures incubated at 37C for 1-3 days for use in preparing frozen glycerol stocks as well as extraction of genomic DNA for sequencing. Sterile tissues (liver and spleen) that were collected from the same mice using a second set of dissection instruments were processed in parallel, in order to recover contaminants of the bacterial isolation procedures [referred to as “background” (B) strains].

### Genome Sequencing and Analysis

Extraction of high molecular weight DNA from overnight cultures of CPs was accomplished using the Nanobind CBB kit (PacBio catalog no.102-301-900). Barcoded sequencing libraries were prepared using the SMRTbell prep kit 3.0 (PacBio catalog no.102-141-700) with SMRTbell barcoded adapter 3.0 (PacBio catalog no.102-009-200) and using the Covaris g-TUBE shearing method to generate DNA fragments of 7-12 kb in length. Quantification of DNA concentrations was accomplished using the Qubit fluorometer method. The barcoded libraries were sequenced using a PacBio Sequel IIe system with HiFi workflows; one SMRT Cell 8M was used for run times of 15 hrs.

Raw sequencing reads were assembled using canu v2.2 with its standard options^64^. Resultant contigs were manually inspected, and BLAST was used to identify and remove duplicate assembled contigs. Phylogenetic placement in the GTDB release 202 system used the ANI-based script speciate.pI^65,66^. Phylogenomic analysis (Figure 1A) was performed using Species Tree Builder in KBase^67^. Genome assemblies were submitted to NCBI; assembly IDs are forthcoming.

### Comparison of CP *versus* Airway Microbiome Phylogenetic Assignments

Sequencing reads from 4 independent 16S rRNA profiling investigations of the mouse airway microbiome^21–25^ were aligned against refseq genome assemblies and placed into taxonomic groups using an established software pipeline called RapTOR. Sequence quality filtering was performed to remove low-quality ends as well as any partial primer sequences, as described previously^68,69^. The Duskmasker software tool was applied in order to remove low complexity and unidentified reads^70^. FASTQ sequences with low quality scores were then removed, followed by removal of any host sequences as identified through alignment against the mouse genome sequence (GRCm38) using Bowtie 2^71^. An additional precise alignment with mitochondrial and ribosomal genome sequences was performed to remove any of these host sequences. The remaining non-host (unaligned) sequences were then aligned to the Silva SSU database^72^ using Bowtie 2 with local alignment settings. The generated “.sam” files were converted to “.taxsum” files using SAMtools^73^ in a custom Perl script for each sample, producing a table for read abundance at each taxonomy level, with reads aligning equally well to multiple Silva entries placed through application of the lowest common ancestor (LCA) algorithm. Finally, reads mapping to each taxonomic group represented in a given profiling dataset were enumerated; and each group was assigned a rank based on the relative abundance of reads mapping to it (i.e., the prominence with which it was represented in the profiling dataset). This information is summarized for the taxonomic groups to which the CPs [and, for comparison, the B strains (contaminants)] belong (Tables S1-3).

### Lung Simulating Medium (LSM) Preparation

LSM was made according to Palmer (2006)^17^. Adjustments were made to the original formulation to increase sodium and chloride concentrations to more closely match those in healthy lungs. As a result, LSM contains 92mM NaCl, instead of 66.6mM NaCl as in SCFM^74^. Similarly, glucose concentration was increased to 5.5mM.

### Competition Assays

3ml monocultures of individual CPs and *Bt* were inoculated from frozen glycerol stocks and grown in LSM at 37C with shaking (278rpm) for 24 hours. Cultures were then serially diluted again in LSM at a 1:50 ratio, and grown at 37C with shaking until late log phase, at which point the cultures were centrifuged at 3000g for 10 min and resuspended in fresh LSM. CPs were then diluted according to the ratios being tested, where a 1:1 ratio represented a cell suspension of 3.33×10^4^ CFU/mL, a 0.1 ratio represented 3.33×10^3^ CFU/mL, etc. After dilution, 3.33×10^4^ CFU/ml of *Bt* was added to each 15ml CP dilution tube as well as to a *Bt*-only control. The 15ml volume was split into 5 3ml aliquots to make biological replicate assays. All samples were incubated at 37C with shaking for 24 h.

After 24 hours, samples were removed, diluted 1:100-1:100,000 in PBS, and plated onto LB plus carbenicillin (100μg/mL) to select for *Bt* colonies. Plates were incubated at 37C for 48 h or until colonies became visible, and colony forming units (CFU) were enumerated. Percent *Bt* growth was calculated by taking the CFU/ml of each replicate culture and dividing by the average of the CFU/ml in the *Bt*-only control and multiplying by 100. Dose-response curves and analyses were generated using the ECAnything algorithm in Graphpad Prism 10.

### Competition Assays Measuring CP Cell Densities

As the CPs were generally sensitive to antibiotics, it was not possible to measure their abundance in competition assays using the antibiotic selection plating method outlined in “Competition Assays”. Monocultures of each CP and Bt were inoculated from frozen glycerol stocks and grown in LSM at 37°C with shaking (220rpm) overnight. Cell density of the monocultures was measured using a Quantom Tx Microbial Cell Counter.

Cultures were diluted to 3.33E4 CFU/ml in LSM, and 800ul of each CP were loaded into 6 chambers of the Cerillo Co-Culture Duet System (cat#NC2389319). The opposite side of 3 chambers was loaded with 800ul of Bt in LSM, or of LSM alone as a negative control. Plates were incubated with shaking at 350rpm at 37°C for 24 hours, and then 100ul of culture were removed from the CP side of each chamber and diluted 1:10. The diluted samples were counted using the Quantom Tx Microbial Cell Counter.

### Phage Isolation and Plaque Assays

A 3ml culture of each CP was grown at 37C with shaking (278rpm) for 16 h, at which point 2ul of the culture were added to a well of a 2ml deep well plate containing 1ml of LB. These cultures were grown at 30C with shaking (750rpm) for 24 h, at which point each culture was back-diluted 1:70 and grown at 30C with shaking until it reached an OD of 1. To each of these 1ml cultures, 2ul of 3% hydrogen peroxide were added in order to induce phage production. After incubation at 30C with shaking for 16 h, the cultures were centrifuged to pellet cells, and the supernatant was collected and sterilized using a 0.22uM filter. Ammonium sulfate was added to 30% saturation, and precipitate was collected by centrifugation at 16000g for 40 minutes at 4C. The supernatant was decanted, and the phage-containing precipitate was resuspended in 1ml of SM buffer (100mM NaCl, 8mM MgSO_4_•7H_2_O, 50mM Tris-Cl, 0.01% gelatin) and stored at 4°C for use in plaque assays.

To perform plaque assays, 3ml of Bt was grown at 37C with shaking (278rpm) for 16 h, at which point it had reached stationary phase. Standard LB agar was prepared in 100×15mm Petri dishes (BD Biosciences) and allowed to cool. To create the soft agar overlay, 200ul of the Bt culture were mixed with 5ml molten LB soft agar (0.5%) and spread over the standard LB agar in each plate. The plates were allowed to cool for 20 min. Phage-containing precipitate was serially diluted 1:10, and 2ul of the solution was pipetted onto the soft agar and allowed to dry for 10 min. The plates were then incubated at 37C for 24 h, and the presence or absence of plaques was observed.

### Supernatant Inhibition Experiments

CPs and *Bt* were grown in LSM monocultures as previously described. The cultures were diluted to 5×10^5^ CFU/ml, and each CP was combined with *Bt* in co-cultures; additionally, a *Bt*-only control was generated. The co-cultures were grown at 37C with shaking (278rpm) for ∼72 h, centrifuged to pellet cells, and the supernatants collected and sterilized using a 0.22uM filter. The sterilized supernatant samples were pH adjusted to 7.03. Each supernatant sample was mixed 1:1 with 2X-concentrated LSM, and exponential-phase *Bt* was added to each mixture at 3.33×10^4^ CFU/mL. Growth curves were performed using a Tecan Spark plate reader, measuring OD600 every 0.25 h during growth at 37C with shaking (240rpm) for 30 h. Fold-change in maximum Bt CFU/ml reached was calculated by dividing the maximum CFU of Bt grown in supernatant from itself divided by the maximum CFU of Bt grown in supernatant from each CP co-culture.

### Knockout of *sfp* in CP8

The CP8Δ*sfp* mutant was generated using a double-crossover homologous recombination method described previously^75^. Briefly, homologous arms were amplified up- and down-stream of the *sfp* gene using PCR, and the amplicon was cloned into the T2(2)-ori plasmid in addition to the oriT/traJ region which facilitates conjugative transfer using Golden Gate assembly. The resulting plasmid was introduced into CP8 using an established method for conjugation in *Bacillus*^76^. The transconjugants were plated onto LB supplemented with kanamycin (20 μg/mL) and polymyxin B (5 μg/mL) to select for plasmid-bearing CP8. Colonies recovered from the selection plates were used to inoculate 3ml LB plus kanamycin (20 μg/mL), and the resulting cultures were grown at 37C with shaking (278rpm) for 16 h. 0.05 ml of each culture were then spread on an LB plus kanamycin (20 μg/mL) plate that was incubated at 45C for 16 h in order to induce the first crossover event. A single colony positive for the first crossover event by PCR was recovered from each plate and passaged 6 times in LB liquid culture. Liquid cultures were streaked onto 10 LB agar plates to generate single colonies. Replica plating of 96 colonies was performed to check for loss of kanamycin resistance, and PCR was used to screen for colonies in which *sfp* was absent. Resulting fragments and the 16S region were sequenced to confirm deletion and strain identity.

### Growth Curves of CP8 and the CP8Δ*sfp* Mutant

CP8 and the CP8Δ*sfp* mutant were streaked on LB agar and incubated at 37C for 16 h. Single colonies were used to inoculate 3ml LB cultures in 14ml plastic tubes, and the cultures incubated at 37C with shaking (278rpm) until they reached early stationary phase. The cultures were then back-diluted to an OD of 0.1 in LB, and 100ul of the cell suspension were added to each well of a 96-well plate, generating 4 replicate assays per condition. Growth curves were performed using a Tecan Spark plate reader, measuring OD600 every 0.16 h during growth at 37C with shaking (240rpm) for 20 h.

### Agar Diffusion Assays

Overnight cultures of *Bt* and CP8 were made in LSM as previously described. *Bt* was diluted to OD 0.5 and streaked onto LSM agar using a sterile cotton swab. Plates were briefly allowed to dry and 2ul spots of CP8 overnight culture were added. Plates were incubated at 37C and checked for the appearance of a zone of inhibition after 48 h.

### Carbon Source Screening

Overnight cultures of CPs and *Bt* in LSM were generated as previously described. Cultures were centrifuged and the pelleted bacteria washed twice with 2x LSM with carbon sources removed. The bacteria were then diluted to an OD of 0.04, and 20 ul were added to each well of a 384-well plate in which individual carbon source from LSM were deposited to produce a final concentration of 10mM. Initial OD600 readings were taken using a plate reader, and then the plates were incubated at 30C with shaking (800rpm) for 3 days before a final OD600 reading was taken. Each CP or *Bt* was considered able to utilize a given carbon source if the final OD600 reading was greater than that of the inoculated blank plus 2 standard deviations.

### Exometabolomics

Overnight cultures of CP7, CP8, CP19 and *Bt* were generated as previously described. The cultures were back-diluted 1:50 and then grown at 37C with shaking (278rpm) to late log phase. 3ml co-cultures were made in LSM combining *Bt* at 3.33×10^4^ CFU/ml with the relevant CP at its IC90. The co-cultures were incubated at 37C with shaking, and 1ml samples were collected at 24 h and 36 h. For collection, each sample was centrifuged at 3000rcf for 10 min, and the . supernatant was removed and syringe filtered to remove residual cellular material. Samples were analyzed via GC-TOF by the West Coast Metabolomics Center (Davis, CA). Data were SERRF normalized to remove batch effects, then normalized to the amount injected. Metabolite concentrations were normalized to a media-only control. For analysis, concentrations of each metabolite were converted to metabolite-contributed electrons using the oxidation-reduction half reaction for each metabolite analyzed. For carbon subset analysis, total electrons from lactate, proline, and aspartic acid in the co-culture supernatant at 24 hours were added. Electron concentrations below 0 indicate that a given metabolite is more abundant in co-culture than the media control.

### Pairwise Combination Competition Assays

Pairwise combination competition assays were performed as described above (“Competition Assays”) with the following adjustment. After resuspension in fresh media, the two CPs of interest were mixed together at a 1:1 ratio to generate the desired final CFU/ml (eg. for a 1:1 ratio with *Bt*, the two CPs were combined to generate a final concentration of 3.33×10^4^ CFU/ml). Subsequent dilutions were made and carried out as previously described.

### Quantification of CP8, CP19 and *Bt* Growth in Co-Culture

qPCR primers and probes for the 16S rRNA region for CP8, CP19 and *Bt* were purchased from IDT with the following sequences:

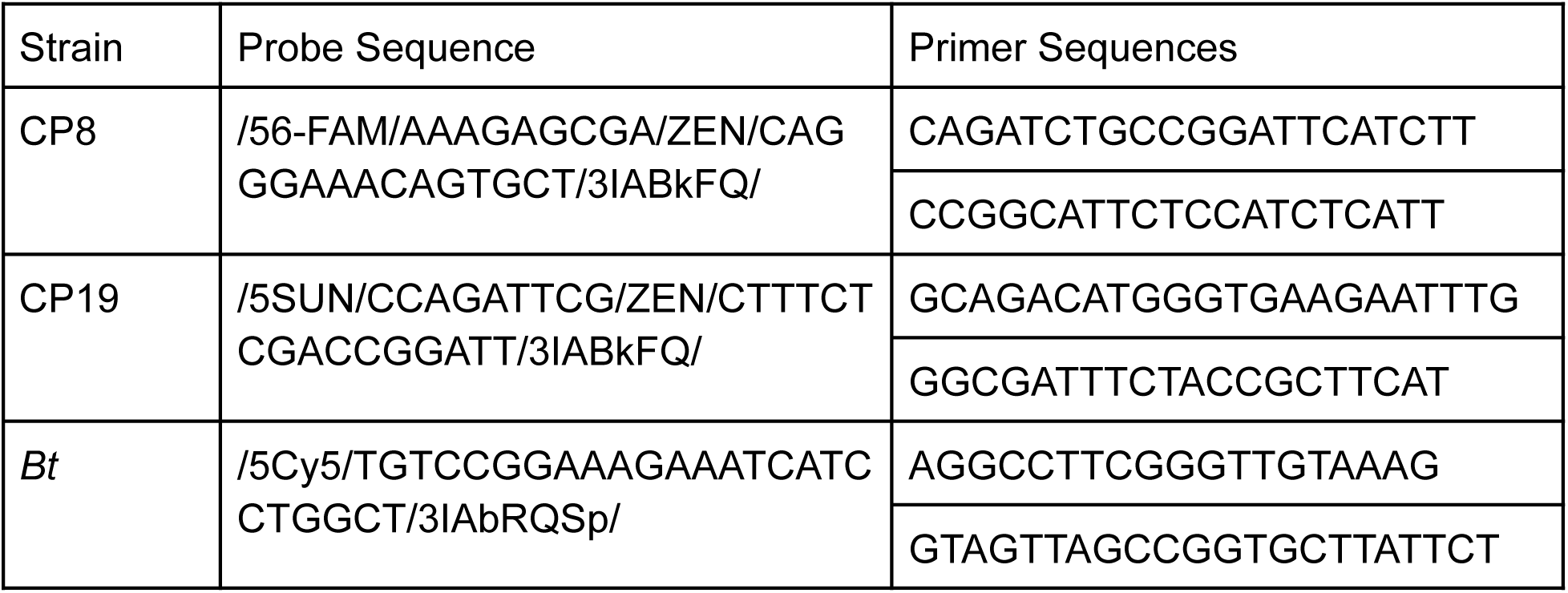

To generate standard curves, overnight cultures of CP8, CP19 and *Bt* were made as previously described in LB. The following day, cultures were backdiluted 1:250 in LB and 1ml of each culture was collected at 11 timepoints. Optical density at each timepoint was measured (OD600 values ranged from 0.05 to 1.033) and cells were pelleted by centrifugation. Genome preparations were done on each pellet using the QIAgen DNeasy Powersoil kit (cat#47014). Concentrations of genomic DNA were quantified using the Qubit dsDNA Quantification Assay Kit (cat#Q32854). A 1:5 dilution of each sample was made to adjust DNA concentration to 3pg-100ng. PrimeTime Gene Expression Master Mix (cat#1055772) was used to run the qPCR reactions in triplicate, with 2ul of diluted genome preparation per well. Samples were analyzed using the BioRad CFX96 Touch Real-Time PCR detection system, with annealing at 60C. To create the standard curve, OD values were plotted against Cq values and a line was fit to the points.

To generate the growth curve, three biological replicates of CP8, CP19 and *Bt* were co-cultured in LSM at 37C and 278rpm for 36 hours, with 1ml collections made every hour between 4-30 hours post-inoculation and an additional collection made at 36 hours. Optical density measurements, DNA extraction and qPCR reaction preparation were performed as previously described. Samples were run in duplicate using the previously established conditions. Measured Cq values from the co-culture samples were converted to OD600 values for each organism using the previously generated standard curve.

### Animal Models

These studies were carried out in strict accordance with the recommendations in the *Guide for the Care and Use of Laboratory Animals* and the National Institute of Health. All efforts were made to minimize suffering of animals. All animals were housed in ABSL2 conditions in an AAALAC-accredited and PHS-assured facility, and the protocol was approved by the Lawrence Livermore National Laboratory Institutional Animal Care and Use Committee (IACUC), which includes ethics in evaluation of protocols. Barrier-housed, specific pathogen-free 6 week old C57Bl/6j mice were acclimated prior to experiments. CP, *Bt*, and animal use were also approved by the Institutional Biosafety Committee (IBC).

### CP Colonization Studies

Each CP was grown in monoculture in LB at 37°C with shaking (250 rpm) for 16 h, recovered through centrifugation, resuspended in PBS, and diluted to an OD600 corresponding to 3.3×10^7^ CFU/mL (based on prior determination of OD600-to-CFU plating efficiency measurements). 30 ul of this dosing material (corresponding to 10^6^ CFU) were administered to the airway (via OPA) of each mouse. After 7 days, the trachea and lungs were collected and homogenized in PBS; dilutions (in PBS) were plated on LB agar; and, after 1-5 days at 37°C, the CFU enumerated in order to assess CP load in the airway tissues.

### Survival Studies

CP dosing material was generated and administered to mice as described above. After 3, 5 or 7 days, the mice were similarly administered (via OPA) 3×10^4^ - 5×10^5^ CFU of *Bt*. Weight, morbidity, and mortality were monitored for 10 days following *Bt* challenge; mice displaying ≥20% loss of body weight were euthanized.

For studies comparing viable versus nonviable CPs, the dosing material was generated as described above but then split into two aliquots, with one remaining untreated (viable CP) and the other subjected to inactivation treatments: Exposure to short-wavelength UV-C light (254 nm) for 20 min in a UV-Box decontamination chamber (Air Science; Fort Meyers, FL) followed by heating at 95°C for 20 min in a dry block incubator (Thermo Fisher Scientific; Waltham, MA). This inactivation protocol causes 100% loss of viability (i.e., no CFU detected after plating on LB) for all CPs tested, as determined through previous experiments and specifically verified for the dosing materials prepared for the studies reported here (data not shown). The viable and nonviable CP dosing materials were then separately administered to mice; after 3, 5, or 7 days the mice were challenged with *Bt* (3×10^4^ - 5×10^5^ CFU); and weight, morbidity, and mortality were monitored for 10 days post-challenge (all as described above).

### Quantification of CP and Pathogen Loads in Airway Tissues

CP dosing material was generated and administered to mice as described above. After 3 days the mice were challenged with *Bt* (3×10^4^ - 5×10^5^ CFU) as described above. After an additional 3 days (i.e., at 3 days post-challenge) the airway tissues were collected and homogenized, and dilutions were plated on LB agar, as described above. After incubation at 37°C for 24 hours the CP CFU were enumerated in order to assess CP load in the airway tissues. After incubation at 37°C for an additional 24-48 hours the *Bt* CFU were enumerated in order to assess *Bt* load in the airway tissues. Note that CP versus *Bt* CFU were easily distinguished based on colony size and morphology.

### *Ex vivo* Lactate Depletion and Pathogen Inhibition Studies

A single colony of CP19 was grown in LSM at 37°C overnight. The culture was centrifuged and the pellet washed twice with PBS followed by resuspension in PBS. Cell density in the suspension was quantified using a Quantom Tx Microbial Cell Counter. The suspension was then backdiluted to 3.3×10^7^ CFU/ml, and 30ul of this diluted suspension, or of PBS (negative control), were added to 120ul of murine lung homogenate. A total of 5 replicate homogenate-based cultures were made per condition, housing them in a 2ml deep-well block. The homogenate-based cultures were incubated at 37°C with shaking at 150rpm for 24 hours, and then centrifuged and sterile filtered. Quantification of lactate levels in the CP19-conditioned and PBS-treated tissue homogenate filtrates was accomplished using the Promega Lactate-Glo Assay (cat#J5021). *Bt* was grown in LSM, centrifuged and washed twice with PBS, and then resuspended in PBS to 5×10^7^ CFU/ml. CP19-conditioned and PBS-treated tissue homogenate filtrates were inoculated with *Bt* (10^6^ CFU) and incubated at 37°C. Growth of the pathogen in the filtrates was monitored by measuring the OD600 every 15 minutes using the Agilent Biotek Synergy Neo2.

### Statistical analyses

Ordinary one-way ANOVA with Tukey’s multiple comparisons test, ROC curve analyses, Mantel-Cox survival analyses, Holm-Šidák multiple comparisons tests and all plotting of data to generate graphs, was completed using Graphpad Prism 10.

### Key Resources

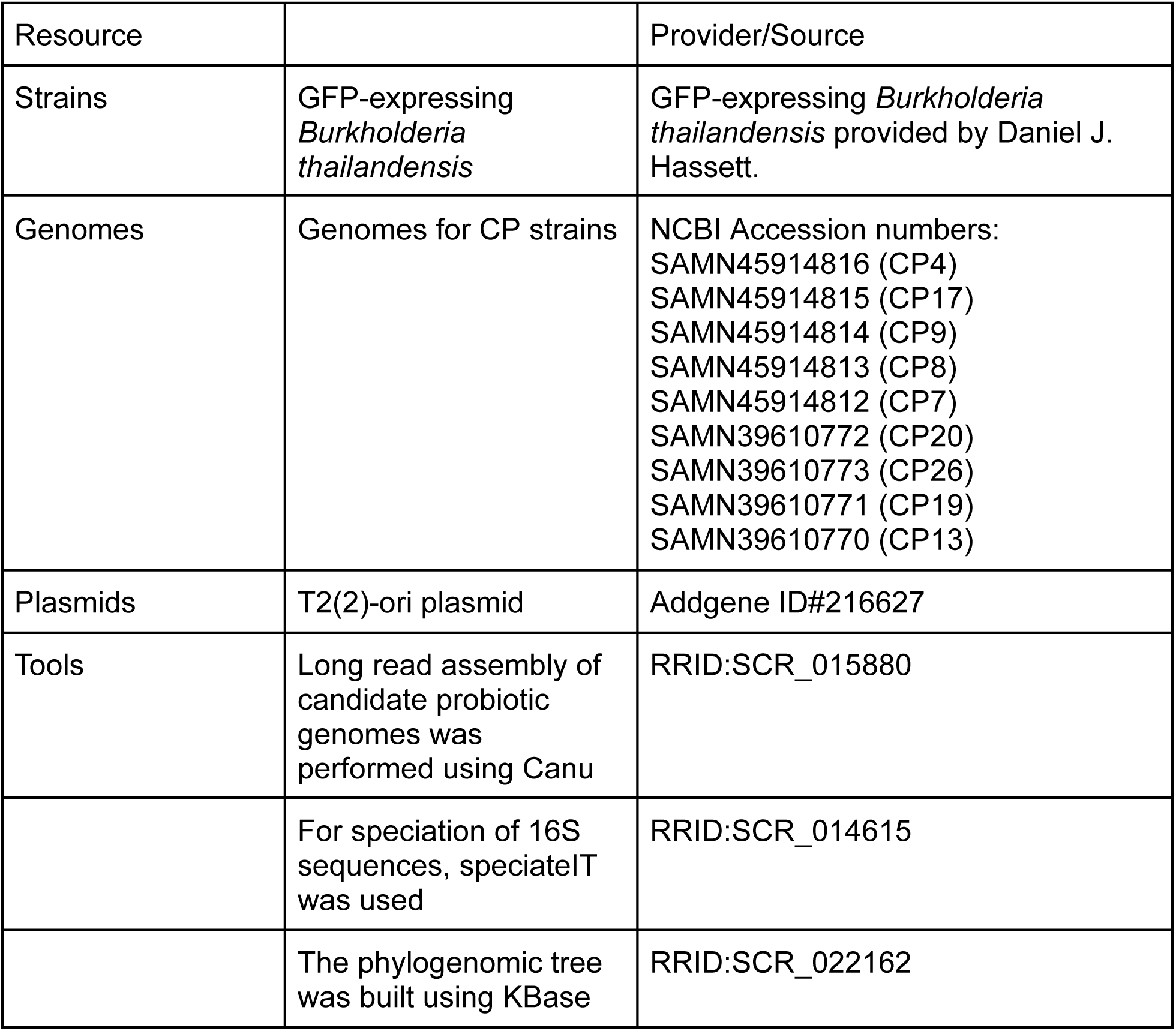

| Resource |  | Provider/Source |
| --- | --- | --- |
| Strains | GFP-expressing *Burkholderia thailandensis* | GFP-expressing *Burkholderia thailandensis* provided by Daniel J. Hassett. |
| Genomes | Genomes for CP strains | NCBI Accession numbers:<br>SAMN45914816 (CP4)<br>SAMN45914815 (CP17)<br>SAMN45914814 (CP9)<br>SAMN45914813 (CP8)<br>SAMN45914812 (CP7)<br>SAMN39610772 (CP20)<br>SAMN39610773 (CP26)<br>SAMN39610771 (CP19)<br>SAMN39610770 (CP13) |
| Plasmids | T2(2)-ori plasmid | Addgene ID#216627 |
| Tools | Long read assembly of candidate probiotic genomes was performed using Canu | RRID:SCR_015880 |
|  | For speciation of 16S sequences, speciateIT was used | RRID:SCR_014615 |
|  | The phylogenomic tree was built using KBase | RRID:SCR_022162 |

## Supplementary Figures

**Figure S1.**
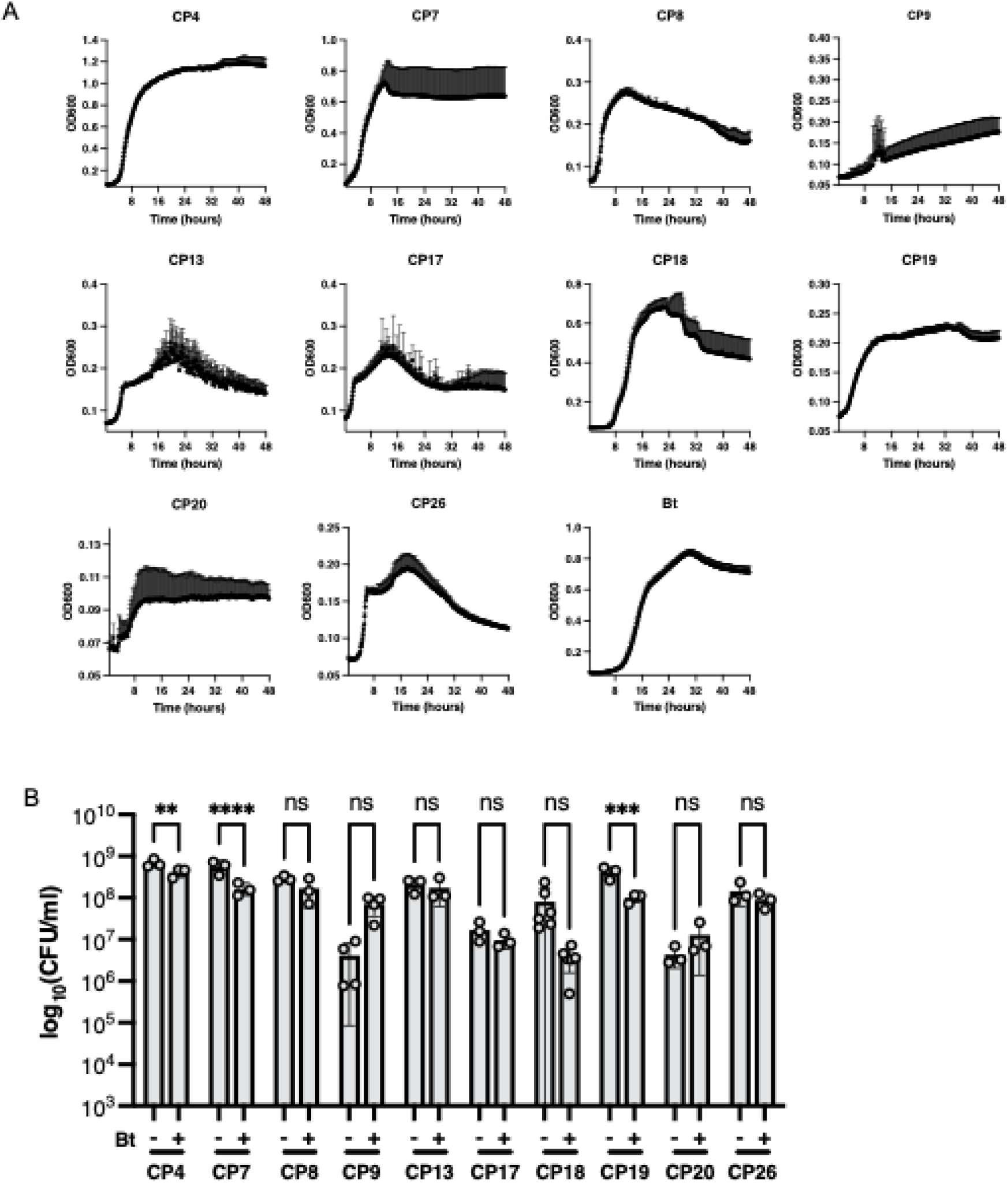
Growth curves in LSM and inhibition of CPs by Bt. (A) CPs and *Bt* were individually inoculated into LSM at 10^6^ CFU/ml, and the cultures were incubated at 37C with shaking (278rpm) for 48 h. (B) Co-culture data measuring CFUs of each CP after co-culture with Bt (+) at a 1:1 ratio or each CP grown alone (-) in LSM. ****P<0.0001; ***P=0.0002; **P=0.0021

**Figure S2.**
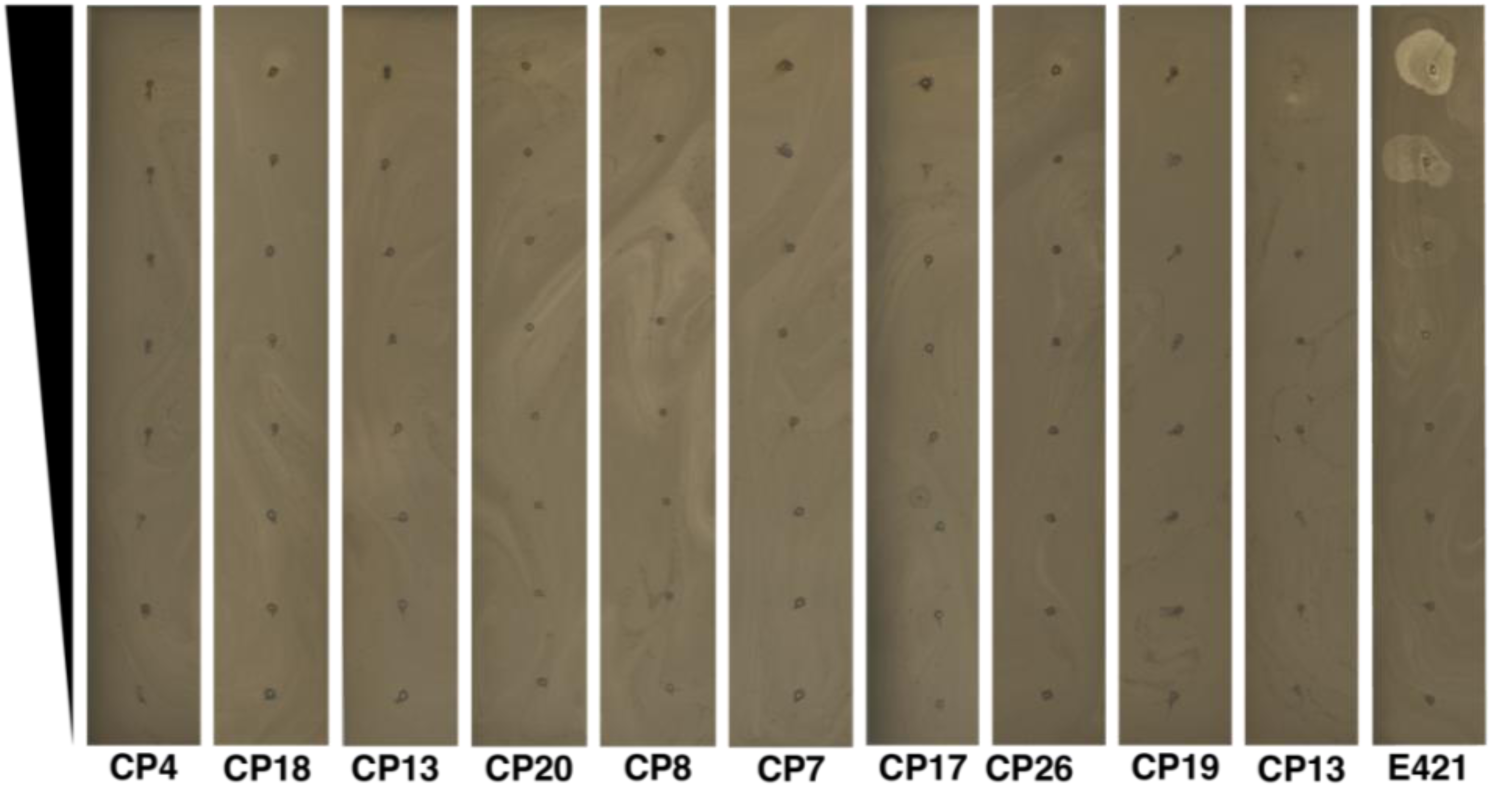
Plaque assays to detect phage isolated from CPs. Supernatants from phage-induced CP cultures were serially diluted 1:10 and spotted onto soft agar containing *Bt*. Supernatant from a phage-induced culture of *Bt* strain E421, which produces a *Bt*-targeting phage, generated visible plaques as expected, demonstrating that the phage detection method was effective.

**Figure S3.**
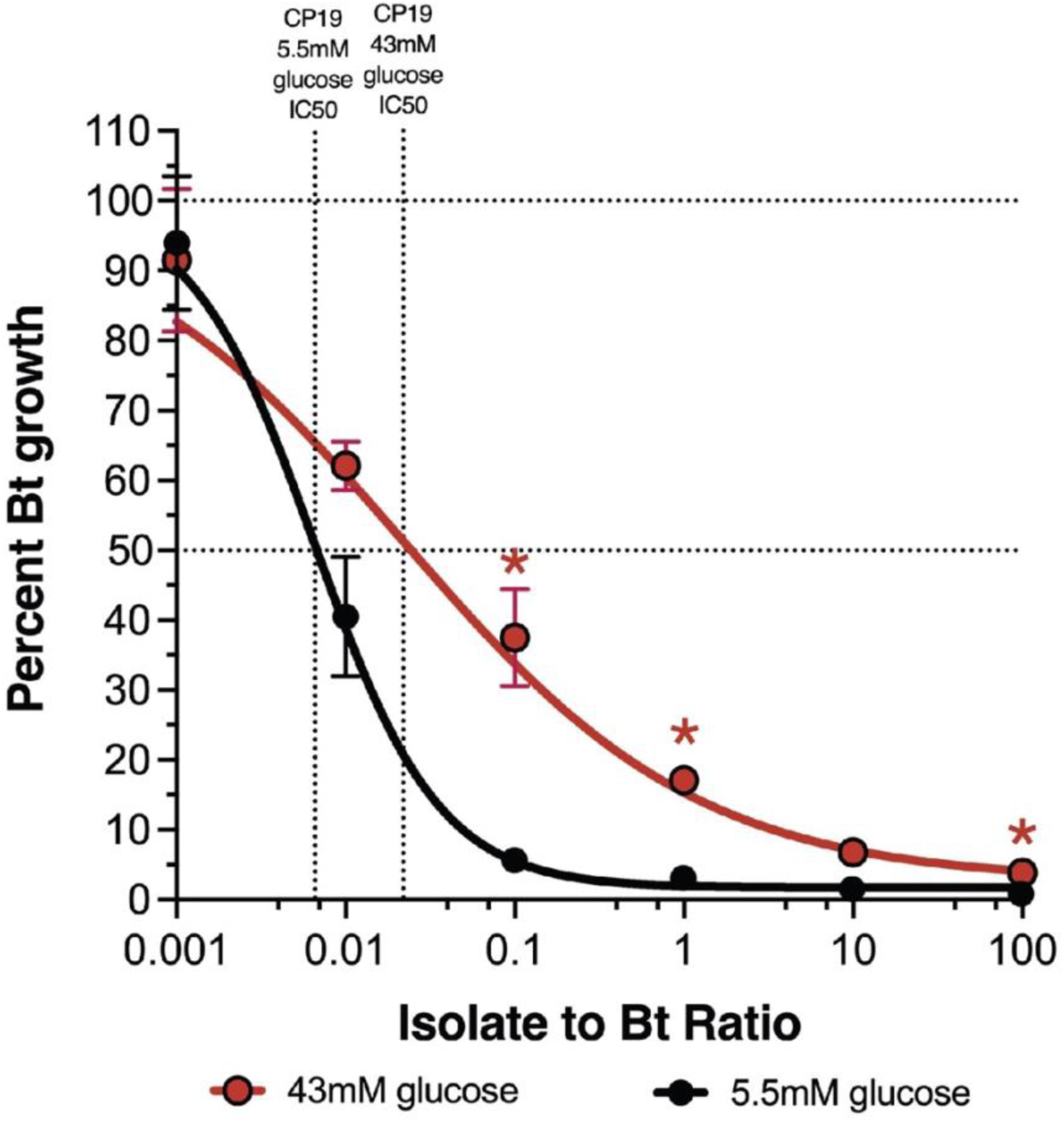
Addition of glucose decreases antagonism between CP19 and Bt. Dose-response curve of *Bt* in co-culture with CP19 in standard LSM (5.5mM glucose) (black) or LSM supplemented to 43mM glucose (red). Vertical dotted lines indicate the IC50 in the low or high glucose condition. Data are represented as the mean ± SEM.

**Figure S4.**
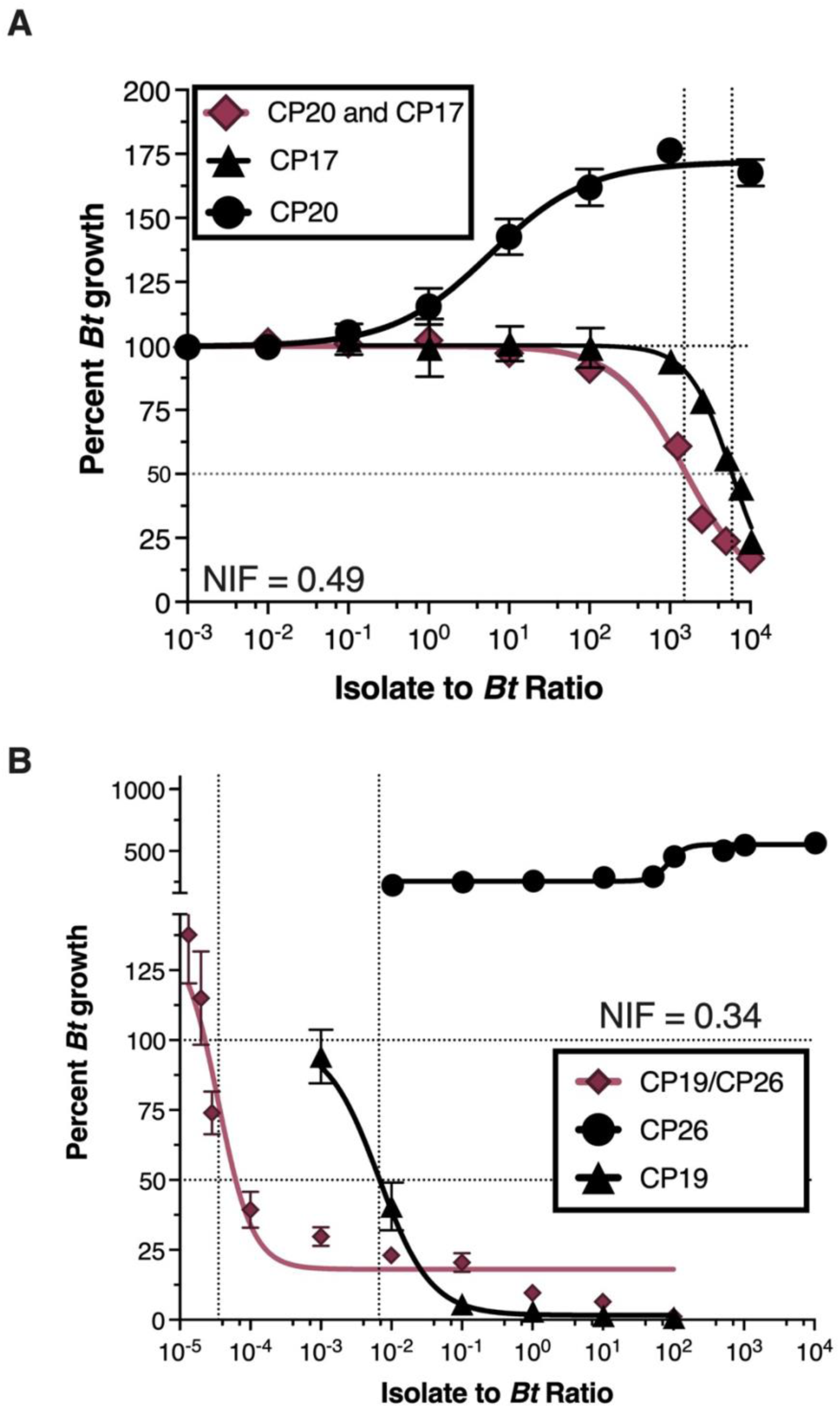
Dose-response curves for pairwise CP combinations with low Niche Index Fractions. (A) IC50 curve for the CP20/CP17 combination (red diamonds). Single strain inhibition curves (black) are shown for CP20 (circles) and CP17 (triangles). (B) IC50 curve for the CP26/CP19 combination (red diamonds). Single strain inhibition curves (black) are shown for CP26 (circles) and CP19 (triangles).

**Figure S5.**
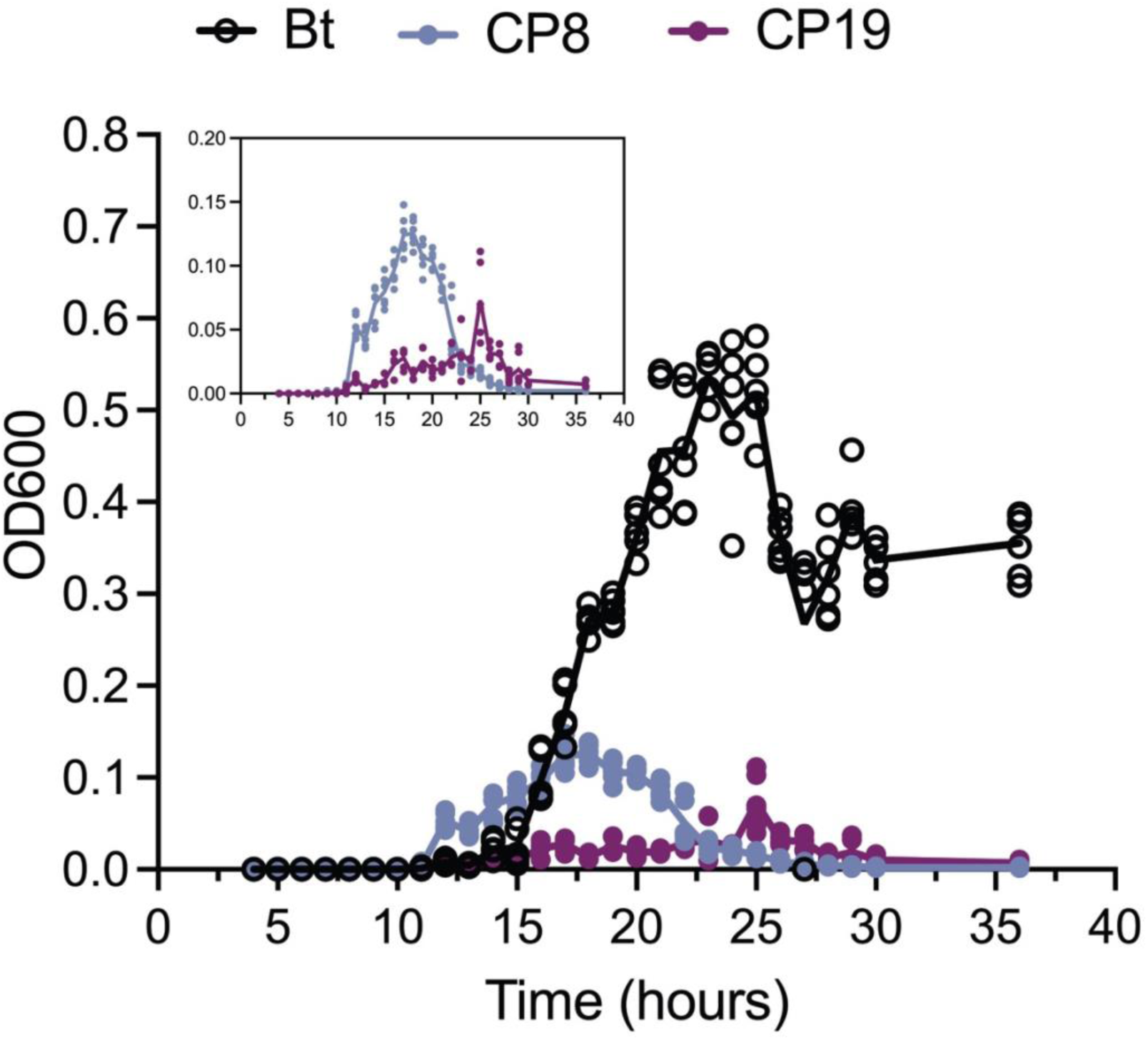
qPCR generated growth curves for CP8, CP19 and *Bt* grown in co-culture. CP8, CP19 and Bt were co-cultured in LSM for 36 hours at 37C and 278rpm. Samples were periodically taken to quantify their OD using qPCR. Three biological replicates of co-cultures were made and two technical replicates of each co-culture analyzed. All 6 points for each organism at each timepoint are displayed. Bt growth curve is denoted as black open circles, CP8 as blue closed circles and CP19 as purple closed circles.

**Figure S6.**
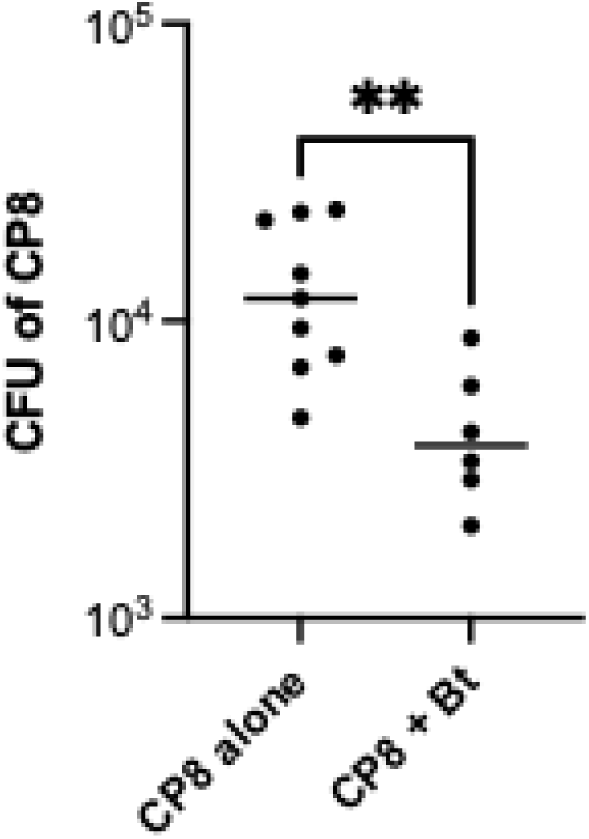
CP8 colonization of the airway is significantly reduced in *Bt*-infected mice. CP8 (10^6^ CFU) was administered to the airway via OPA and after 3 days challenged with either PBS (CP8 alone) or *Bt* (3×10^4^ - 5×10^5^ CFU) (CP8 + Bt) via OPA. Airway tissues were recovered at 3 days post-challenge (6 days post-CP8 administration) for enumeration of CP8 load (expressed as CFU per tissue homogenate). **P<0.002.

**Figure S7.**
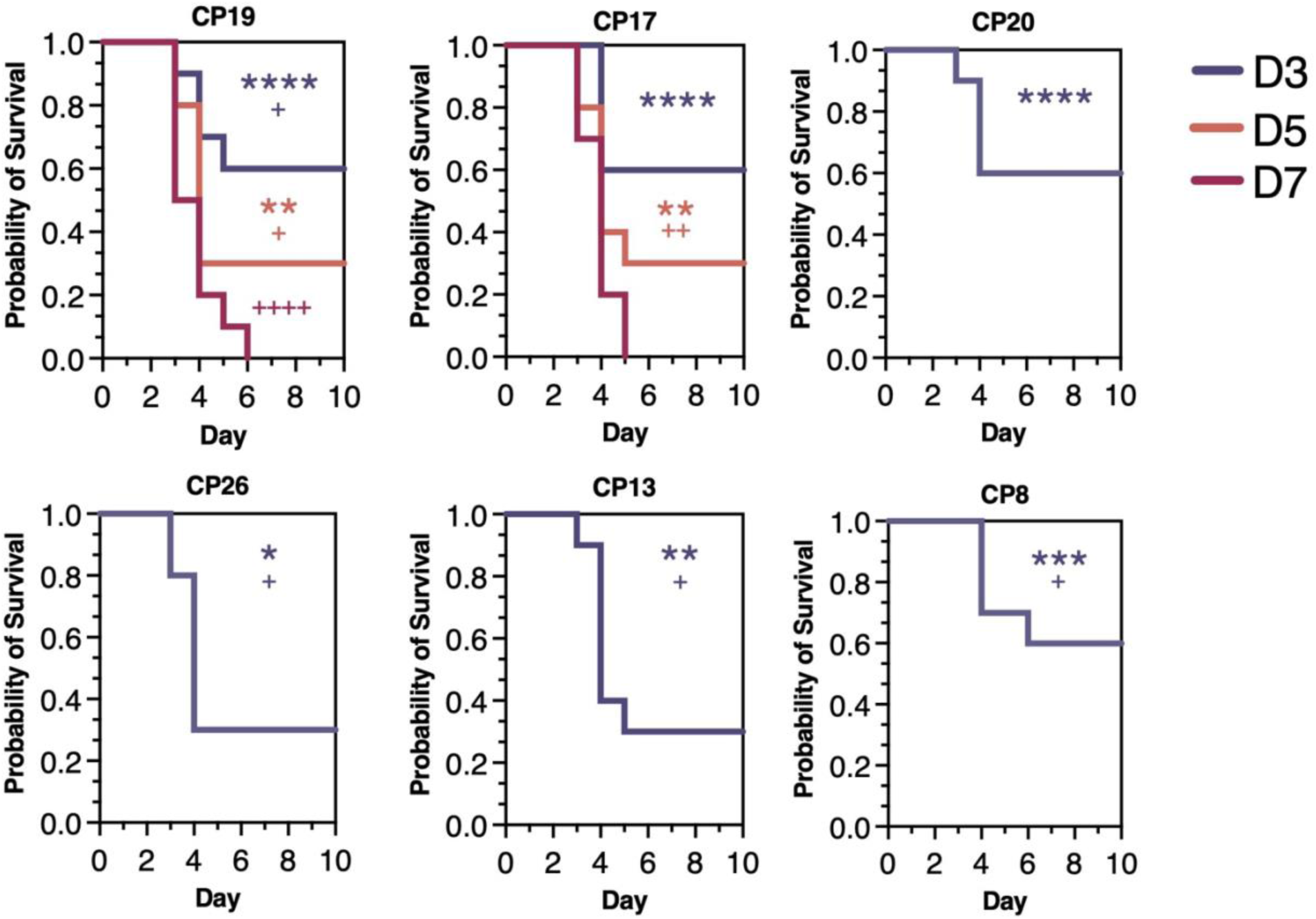
Survival of mice treated with non-viable CPs prior to pathogen challenge. CPs were rendered non-viable (via UV and heat treatment) and administered to the mouse airway (via OPA) at 3 days (blue), 5 days (orange), or 7 days (red) prior to challenge with *Bt* (via OPA). Survival was monitored for 10 days following pathogen challenge. For CP20, CP26, CP13, and CP8 treatment at 3 days prior to pathogen challenge was the only dosing regimen investigated because these CPs provided little to no protection when viable bacteria were administered at 5 days or 7 days prior to pathogen challenge (see Fig. 5E). Asterisks indicate significant differences between survival of mice treated with PBS (negative control) versus the non-viable CP, whereas plus signs indicate significant differences between survival of mice treated with the non-viable CP versus the viable CP, as calculated using the Holm-Šidák multiple comparisons test (****or ^++++^P<0.0001, ***or ^+++^P<0.0002, ** or ^++^P<0.0021, * or ^+^P<0.0332).

**Figure S8.**
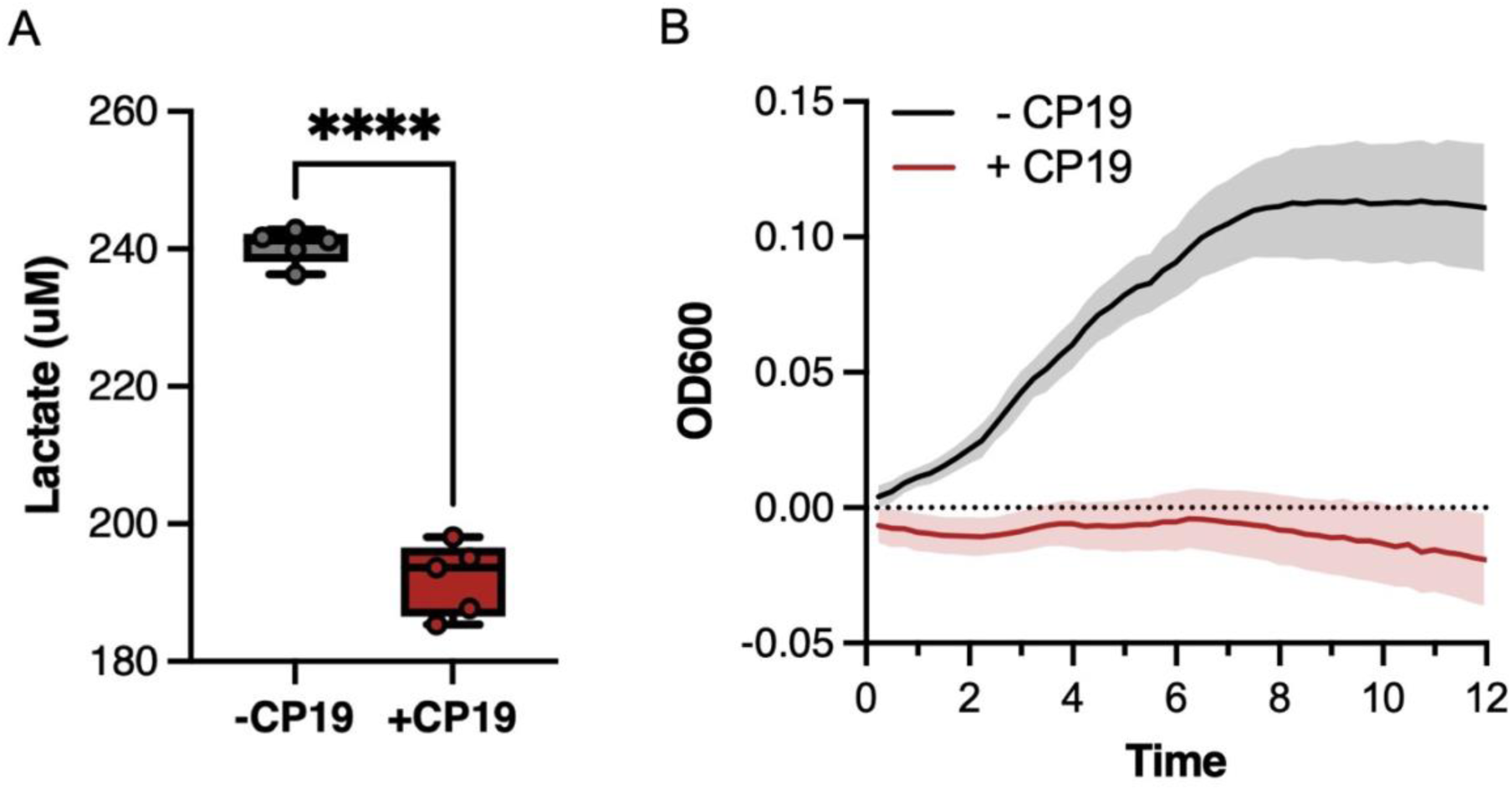
CP19 consumption of lactate and inhibition of pathogen growth in mouse airway tissues *ex vivo*. (A) Lactate levels in airway tissue homogenates treated with PBS (-CP19) or CP19 (10^6^ CFU) (+CP19) for 24 hours. ****P<0.0001. (B) Growth of *Bt* in airway tissue homogenates pre-treated with PBS (-CP19) or CP19 (10^6^ CFU) (+CP19) for 24 hours. Pre-treated tissue homogenates were filter sterilized, the filtrates were inoculated with *Bt* (10^6^ CFU), and OD600 measurements were taken every 15 minutes to monitor growth of the pathogen over the course of 12 hours.

**Table S1.**
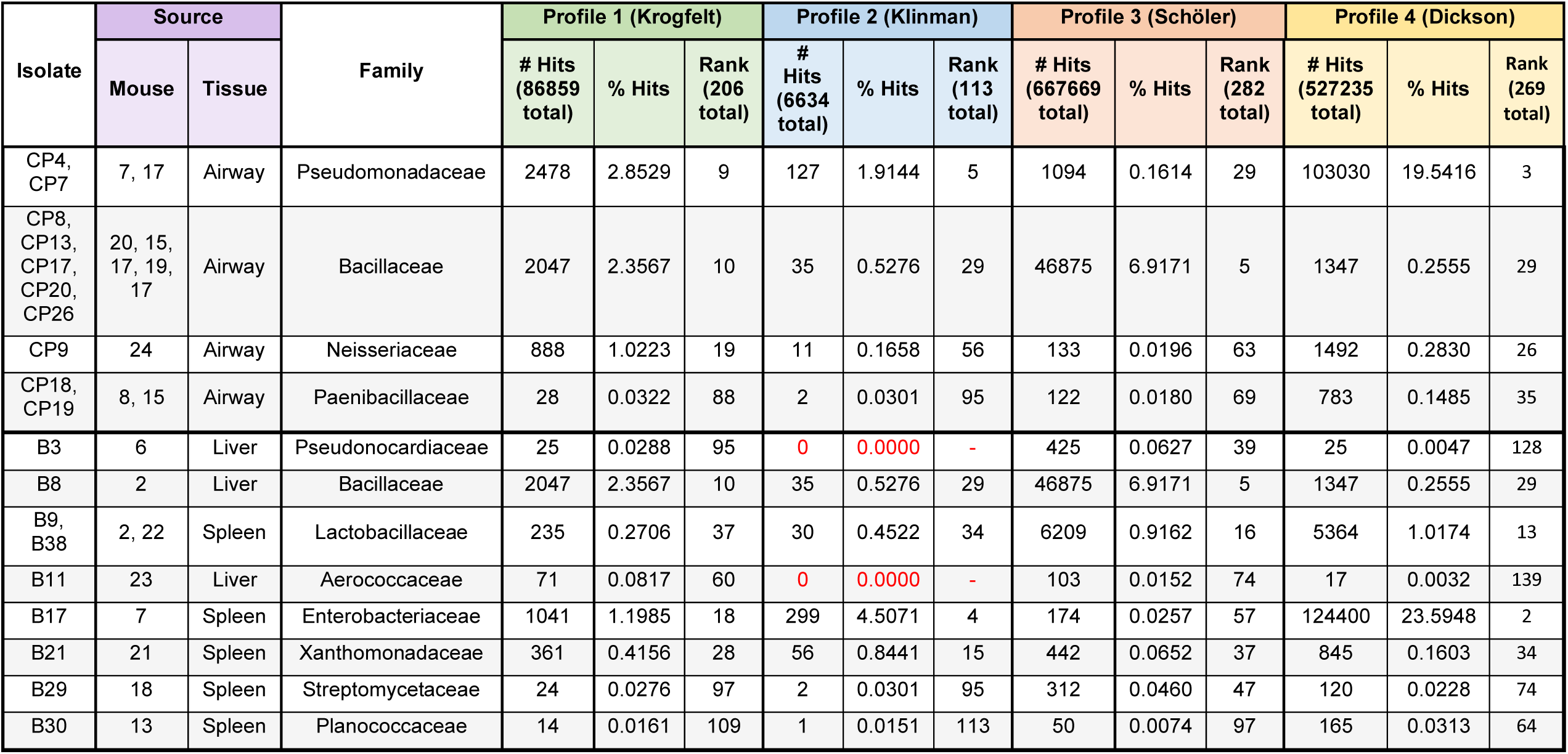
Representation of CP families in mouse airway microbiome profiles. Family-level phylogenetic assignments of the CPs were compared to those of mouse airway microbiome constituents previously identified in four independent 16S rRNA profiling efforts^21–25^. An identical analysis was performed for “background” (B) strains (contaminants recovered from liver and spleen). For each CP and B family, listed are: 1) The number of times that it is represented in a given airway microbiome profile (“# Hits”); 2) Its relative abundance in the profile (“% Hits”); and 3) Its relative rank within the profile, with the most abundant family in the profile designated as “Rank” = 1. For each profile, the total number of 16S rRNA sequences placed at the family level, and the total number of families represented, are listed the headings of the “# Hits” and “Rank” columns, respectively.

**Table S2.**
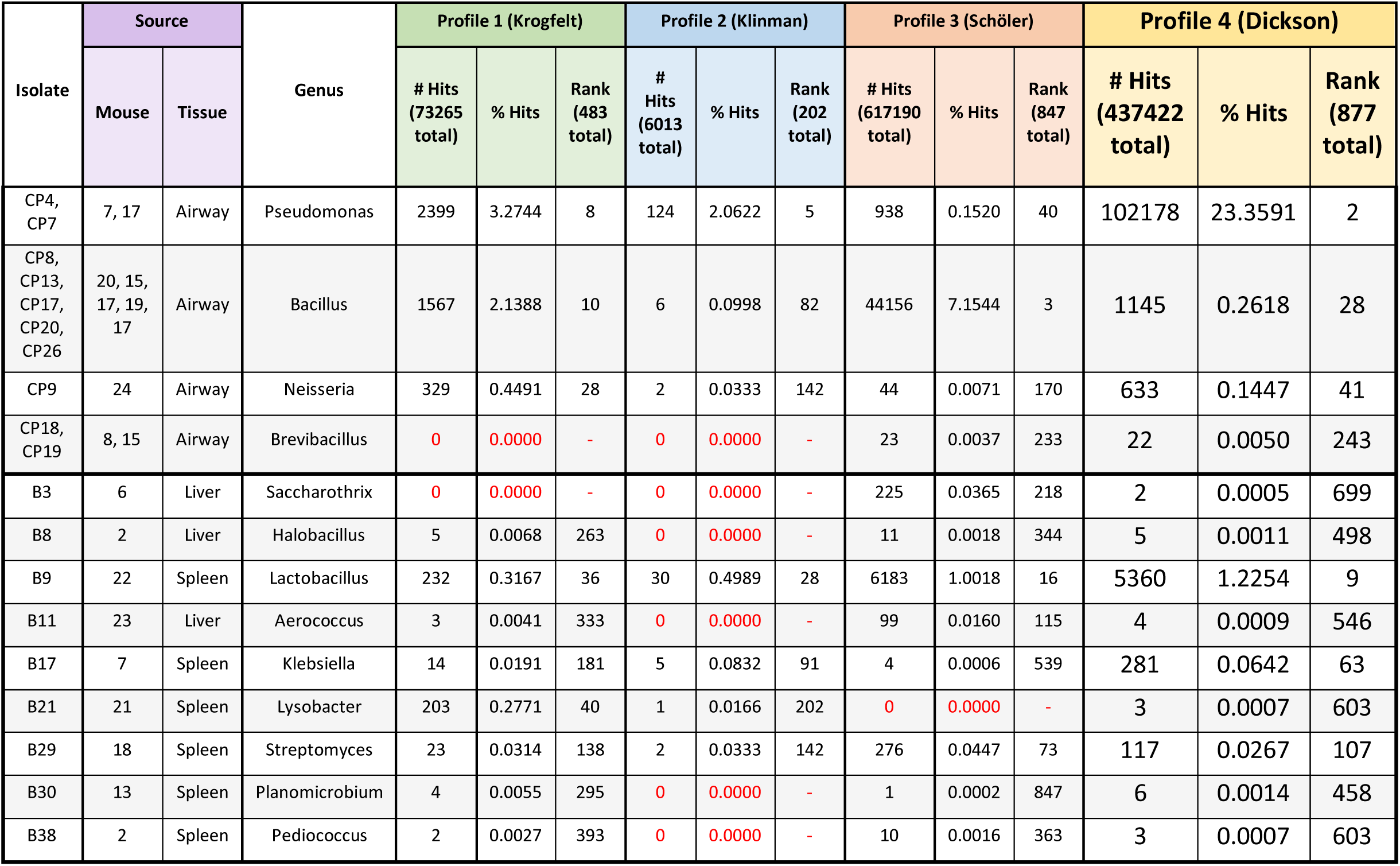
Representation of CP genera in mouse airway microbiome profiles. Genus-level phylogenetic assignments of the CPs, and of the B strains, were compared to those of mouse airway microbiome constituents previously identified in four independent 16S rRNA profiling efforts^21–25^. For each CP and B genus, listed are: 1) The number of times it is represented in a given airway microbiome profile (“# Hits”); 2) Its relative abundance in the profile (“% Hits”); and 3) Its relative rank within the profile, with the most abundant genus in the profile designated as “Rank” = 1. For each profile, the total number of 16S rRNA sequences placed at the genus level, and the total number of genera represented, are listed the headings of the “# Hits” and “Rank” columns, respectively.

**Table S3.**
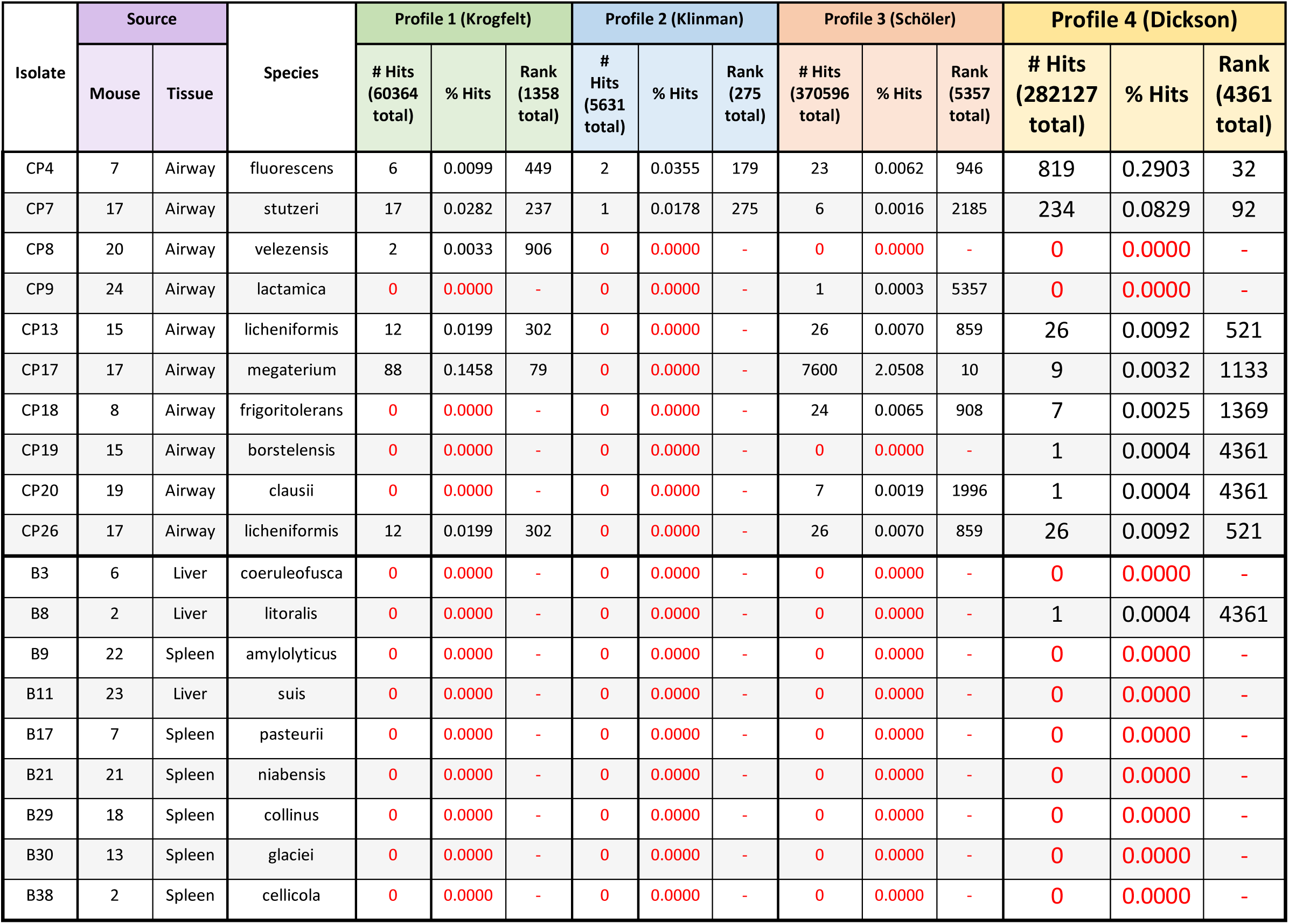
Representation of CP species in mouse airway microbiome profiles. Species-level phylogenetic assignments of the CPs, and of the B strains, were compared to those of mouse airway microbiome constituents previously identified in four independent 16S rRNA profiling efforts^21–25^. For each CP and B species, listed are: 1) The number of times it is represented in a given airway microbiome profile (“# Hits”); 2) Its relative abundance in the profile (“% Hits”); and 3) Its relative rank within the profile, with the most abundant species in the profile designated as “Rank” = 1. For each profile, the total number of 16S rRNA sequences placed at the species level, and the total number of species represented, are listed the headings of the “# Hits” and “Rank” columns, respectively.

**Table S4.**
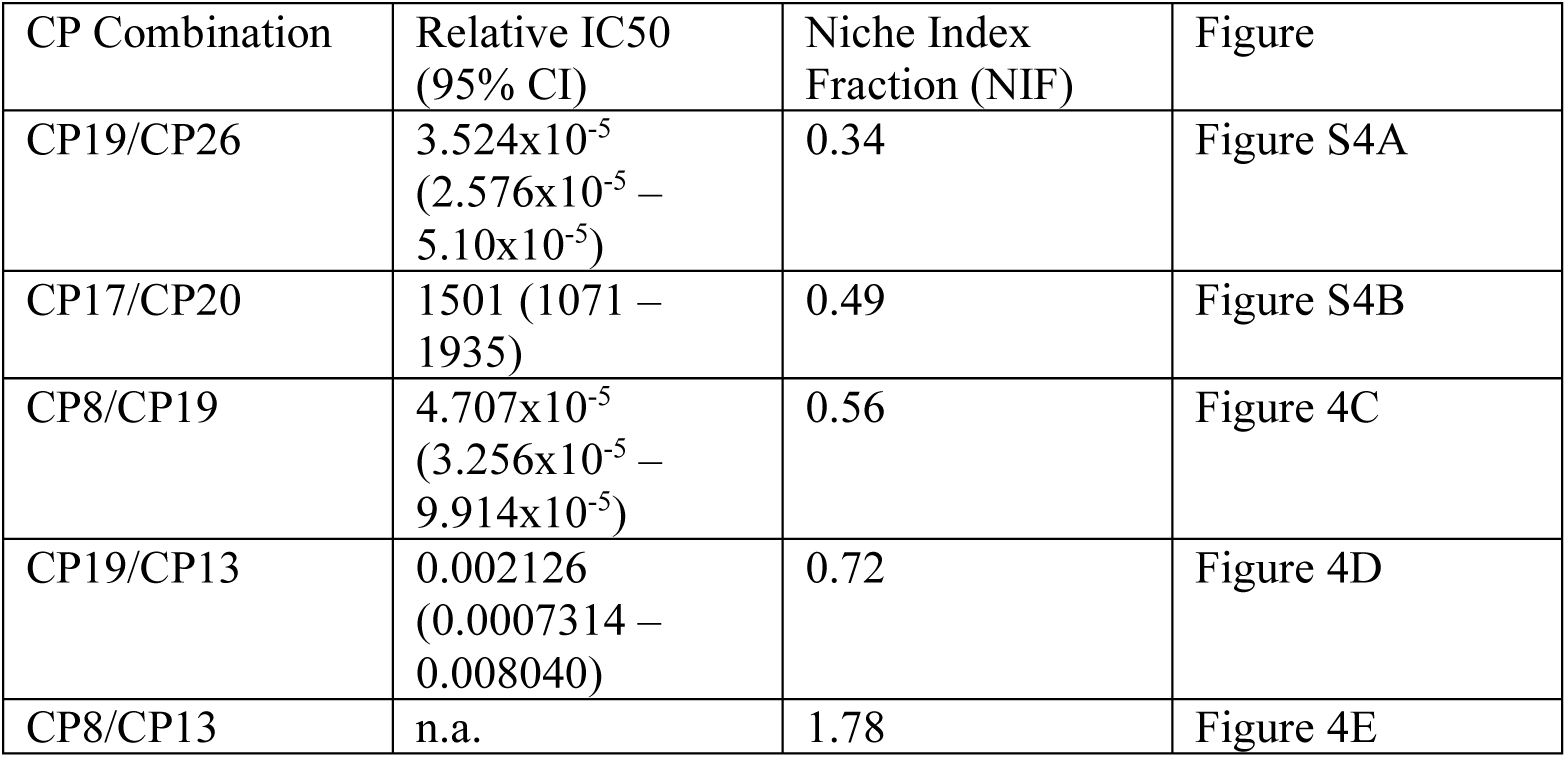
Relative IC50 and Niche Index Fraction values for each CP combination tested. Relative IC50 values are reported as IC50 (95% confidence interval). Non-inhibitory combinations have relative IC50 values listed as n.a. (not applicable).

