## Supplemental Figures for "Niche exclusion of a lung pathogen in mice with designed probiotic communities"

| CP | Species | Relative IC50 | 95% CI | *Bt* growth at a 1:1 ratio (%) | Niche Index |
|---|---|---|---|---|---|
| CP7 | *Pseudomonas stutzeri* | 0.001172 | 0.0006442-0.003346 | 1.893 | 0.875 |
| CP4 | *Pseudomonas fluorescens* | 0.003086 | 0.001791-0.005331 | 3.200 | 0.869 |
| CP19 | *Brevibacillus borstelensis* | 0.006604 | 0.006484 - 0.01141 | 3.046 | 0.775 |
| CP17 | *Bacillus megaterium* | 5738 | 3598 – n.d. | 98.99 | 0.704 |
| CP18 | *Peribacillus frigoritolerans* | 0.0001693 | 0.0001633 - 0.0001850 | 0.534 | 0.591 |
| CP20 | *Bacillus clausii* | n.d. | n.d. | 115.2 | 0.457 |
| CP26 | *Bacillus licheniformis* | n.d. | n.d. | 260.0 | 0.433 |
| CP13 | *Bacillus licheniformis* | n.d. | n.d. | 913.0 | 0.421 |
| CP9 | *Neisseria lactamica* | n.d. | n.d. | 157.1 | 0.357 |
| CP8 | *Bacillus velezensis* | 2.316 | 0.8280 – 5.073 | 56.97 | 0.351 |

**Table 1 Summary of pathogen inhibition by CPs.** Relative IC50 values and 95% confidence intervals (95% CI) are shown for each inhibitory CP. Not determined (n.d.) indicates that a value could not be measured. CPs are listed in order of decreasing NI values.

### Supplementary Figures

A

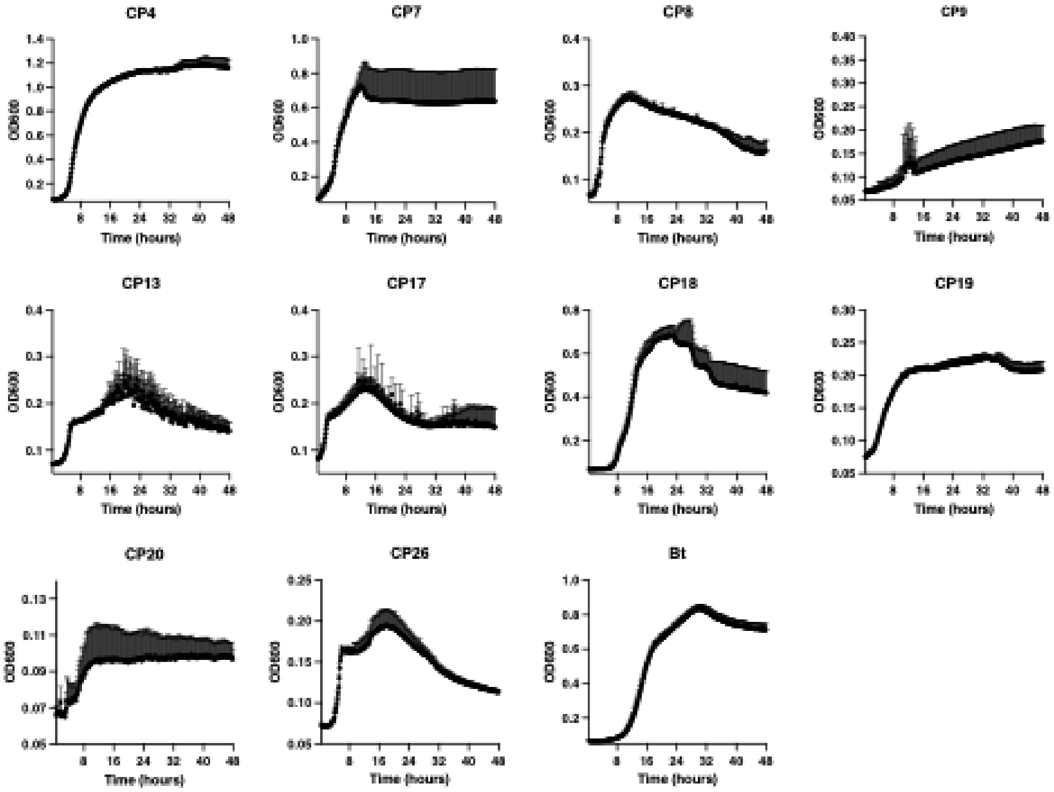


B

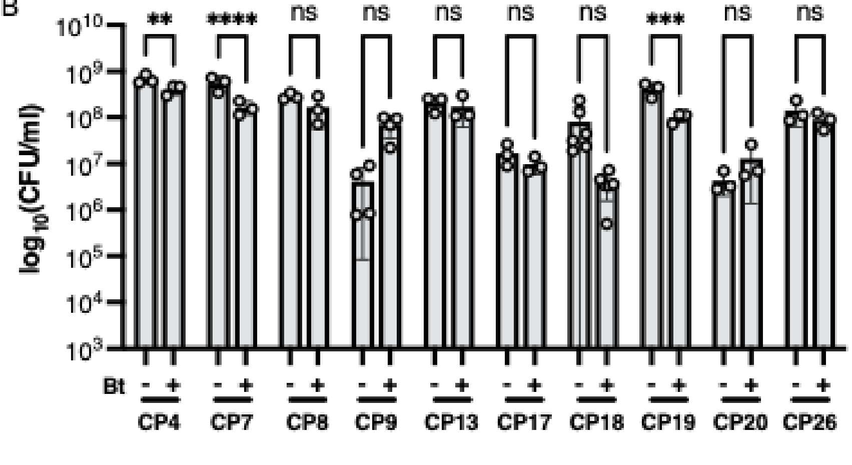


**Figure S1 Growth curves in LSM and inhibition of CPs by Bt.** (A) CPs and *Bt* were individually inoculated into LSM at $10^6$ CFU/ml, and the cultures were incubated at 37C with shaking (278rpm) for 48 h. (B) Co-culture data measuring CFUs of each CP after co-culture with Bt (+) at a 1:1 ratio or each CP grown alone (-) in LSM. ****P<0.0001; ***P=0.0002; **P=0.0021

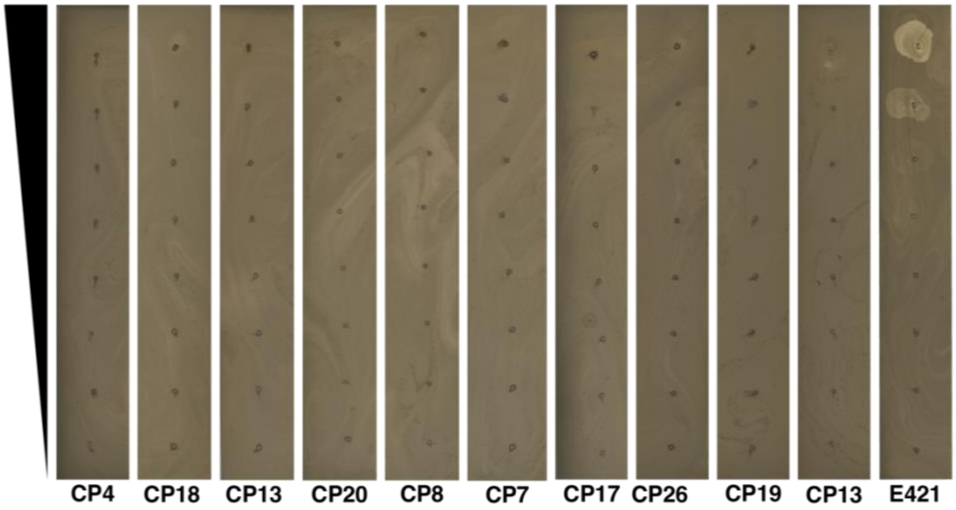


**Figure S2 Plaque assays to detect phage isolated from CPs.** Supernatants from phage-induced CP cultures were serially diluted 1:10 and spotted onto soft agar containing *Bt*. Supernatant from a phage-induced culture of *Bt* strain E421, which produces a *Bt*-targeting phage, generated visible plaques as expected, demonstrating that the phage detection method was effective.

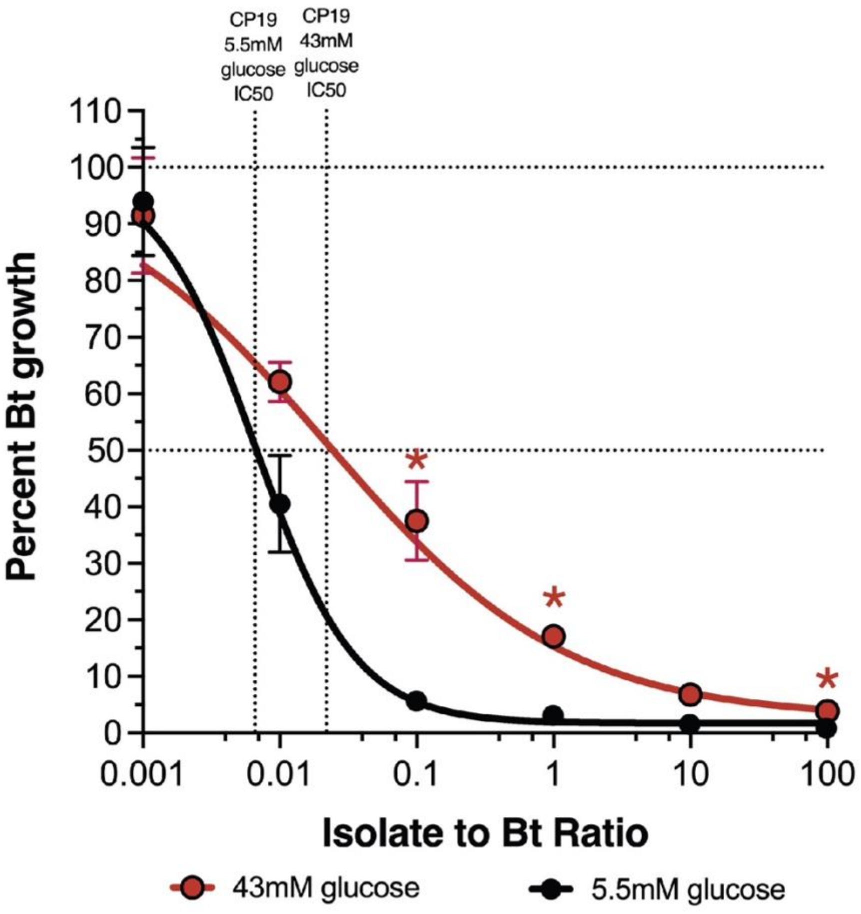


**Figure S3 Addition of glucose decreases antagonism between CP19 and Bt.** Dose-response curve of *Bt* in co-culture with CP19 in standard LSM (5.5mM glucose) (black) or LSM supplemented to 43mM glucose (red). Vertical dotted lines indicate the IC50 in the low or high glucose condition. Data are represented as the mean ± SEM.

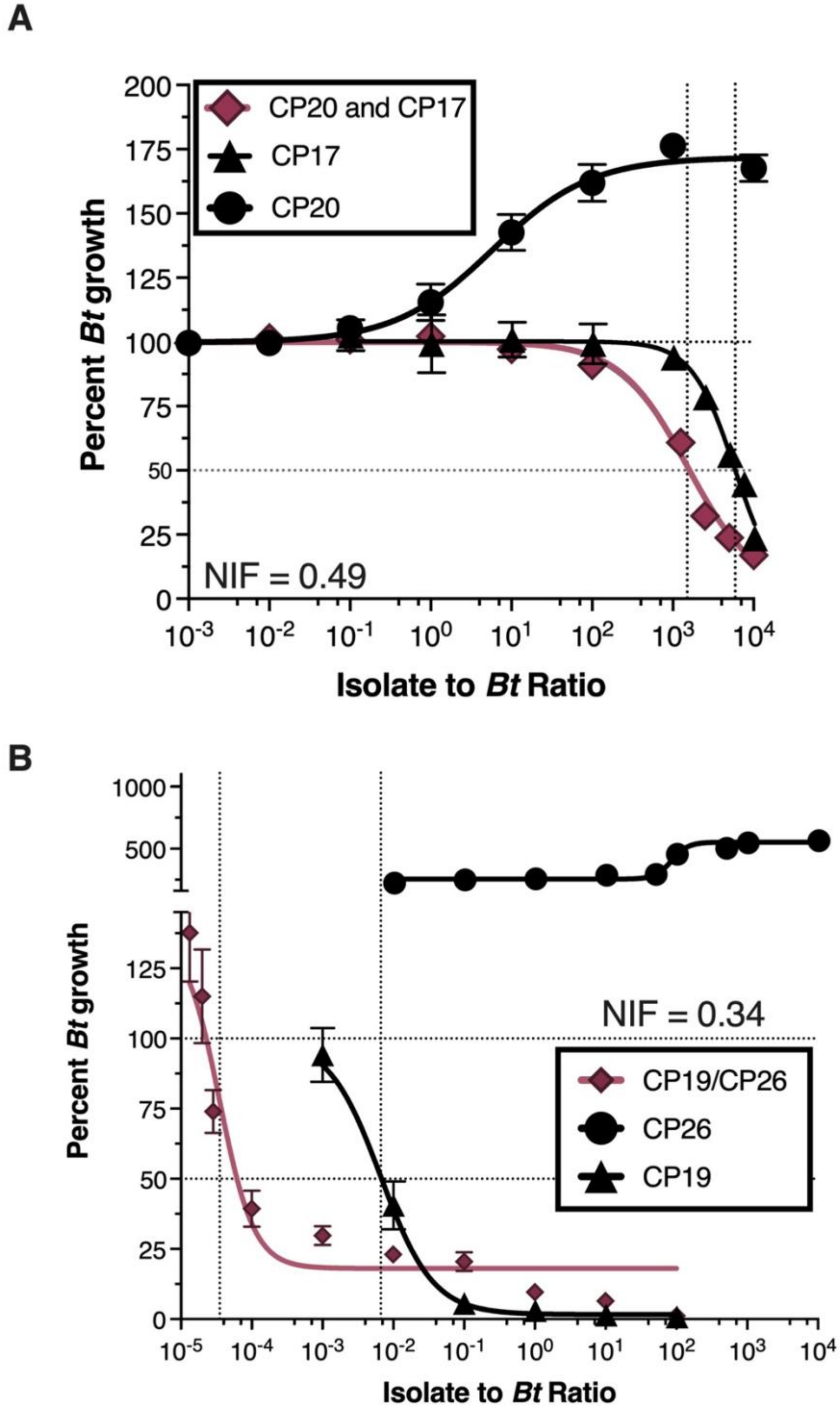


**Figure S4 Dose-response curves for pairwise CP combinations with low Niche Index Fractions** (A) IC50 curve for the CP20/CP17 combination (red diamonds). Single strain inhibition curves (black) are shown for CP20 (circles) and CP17 (triangles). (B) IC50 curve for the CP26/CP19 combination (red diamonds). Single strain inhibition curves (black) are shown for CP26 (circles) and CP19 (triangles).

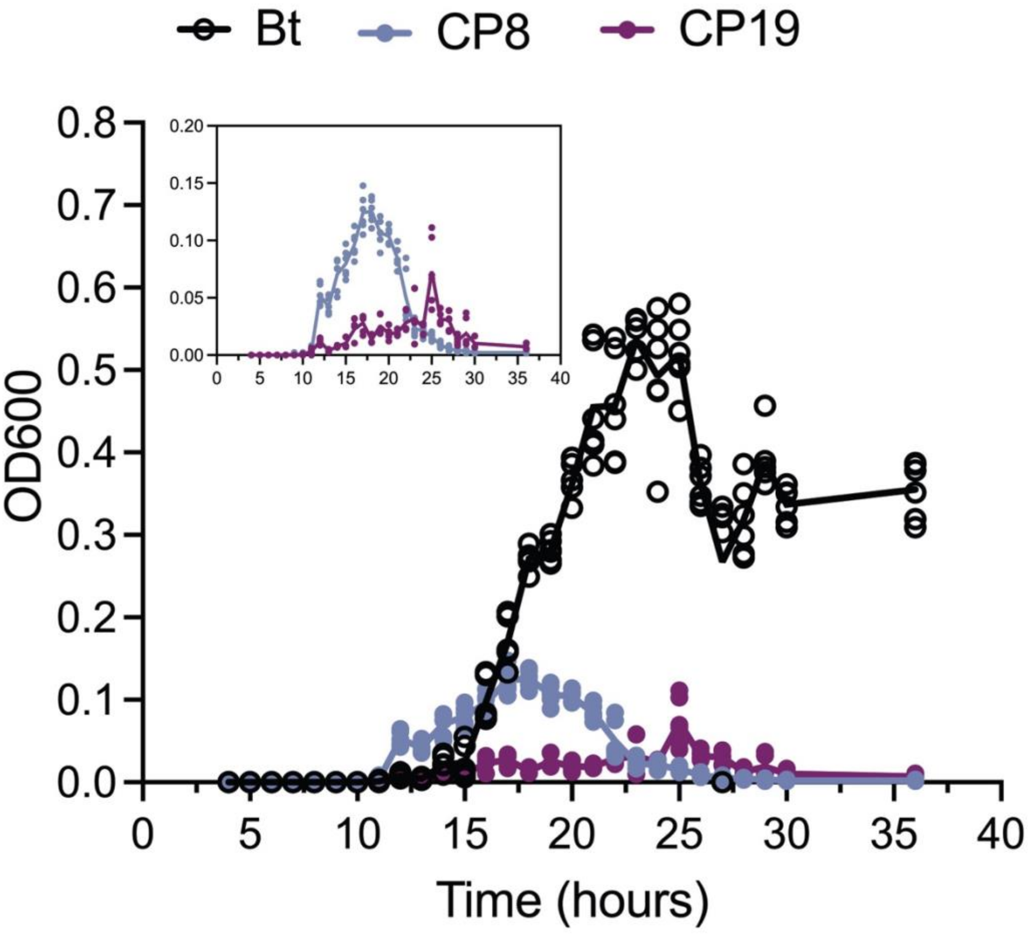


**Figure S5 qPCR generated growth curves for CP8, CP19 and *Bt* grown in co-culture.** CP8, CP19 and Bt were co-cultured in LSM for 36 hours at 37C and 278rpm. Samples were periodically taken to quantify their OD using qPCR. Three biological replicates of co-cultures were made and two technical replicates of each co-culture analyzed. All 6 points for each organism at each timepoint are displayed. Bt growth curve is denoted as black open circles, CP8 as blue closed circles and CP19 as purple closed circles.

| Isolate | Source | | Family | Profile 1 (Krogfelt) | | | Profile 2 (Klinman) | | | Profile 3 (Schöler) | | | Profile 4 (Dickson) | | |
|---|---|---|---|---|---|---|---|---|---|---|---|---|---|---|---|
| | Mouse | Tissue | | # Hits (86859 total) | % Hits | Rank (206 total) | # Hits (6634 total) | % Hits | Rank (113 total) | # Hits (667669 total) | % Hits | Rank (282 total) | # Hits (527235 total) | % Hits | Rank (269 total) |
| CP4, CP7 | 7, 17 | Airway | Pseudomonadaceae | 2478 | 2.8529 | 9 | 127 | 1.9144 | 5 | 1094 | 0.1614 | 29 | 103030 | 19.5416 | 3 |
| CP8, CP13, CP17, CP20, CP26 | 20, 15, 17, 19, 17 | Airway | Bacillaceae | 2047 | 2.3567 | 10 | 35 | 0.5276 | 29 | 46875 | 6.9171 | 5 | 1347 | 0.2555 | 29 |
| CP9 | 24 | Airway | Neisseriaceae | 888 | 1.0223 | 19 | 11 | 0.1658 | 56 | 133 | 0.0196 | 63 | 1492 | 0.2830 | 26 |
| CP18, CP19 | 8, 15 | Airway | Paenibacillaceae | 28 | 0.0322 | 88 | 2 | 0.0301 | 95 | 122 | 0.0180 | 69 | 783 | 0.1485 | 35 |
| B3 | 6 | Liver | Pseudonocardiaceae | 25 | 0.0288 | 95 | 0 | 0.0000 | - | 425 | 0.0627 | 39 | 25 | 0.0047 | 128 |
| B8 | 2 | Liver | Bacillaceae | 2047 | 2.3567 | 10 | 35 | 0.5276 | 29 | 46875 | 6.9171 | 5 | 1347 | 0.2555 | 29 |
| B9, B38 | 2, 22 | Spleen | Lactobacillaceae | 235 | 0.2706 | 37 | 30 | 0.4522 | 34 | 6209 | 0.9162 | 16 | 5364 | 1.0174 | 13 |
| B11 | 23 | Liver | Aerococcaceae | 71 | 0.0817 | 60 | 0 | 0.0000 | - | 103 | 0.0152 | 74 | 17 | 0.0032 | 139 |
| B17 | 7 | Spleen | Enterobacteriaceae | 1041 | 1.1985 | 18 | 299 | 4.5071 | 4 | 174 | 0.0257 | 57 | 124400 | 23.5948 | 2 |
| B21 | 21 | Spleen | Xanthomonadaceae | 361 | 0.4156 | 28 | 56 | 0.8441 | 15 | 442 | 0.0652 | 37 | 845 | 0.1603 | 34 |
| B29 | 18 | Spleen | Streptomycetaceae | 24 | 0.0276 | 97 | 2 | 0.0301 | 95 | 312 | 0.0460 | 47 | 120 | 0.0228 | 74 |
| B30 | 13 | Spleen | Planococcaceae | 14 | 0.0161 | 109 | 1 | 0.0151 | 113 | 50 | 0.0074 | 97 | 165 | 0.0313 | 64 |

**Table S1 Representation of CP families in mouse airway microbiome profiles.** Family-level phylogenetic assignments of the CPs were compared to those of mouse airway microbiome constituents previously identified in four independent 16S rRNA profiling efforts[21–25]. An identical analysis was performed for "background" (B) strains (contaminants recovered from liver and spleen). For each CP and B family, listed are: 1) The number of times that it is represented in a given airway microbiome profile ("# Hits"); 2) Its relative abundance in the profile ("% Hits"); and 3) Its relative rank within the profile, with the most abundant family in the profile designated as "Rank" = 1. For each profile, the total number of 16S rRNA sequences placed at the family level, and the total number of families represented, are listed the headings of the "# Hits" and "Rank" columns, respectively.

| Isolate | Source | | Genus | Profile 1 (Krogfelt) | | | Profile 2 (Klinman) | | | Profile 3 (Schöler) | | | Profile 4 (Dickson) | | |
|---|---|---|---|---|---|---|---|---|---|---|---|---|---|---|---|
| | Mouse | Tissue | | # Hits (73265 total) | % Hits | Rank (483 total) | # Hits (6013 total) | % Hits | Rank (202 total) | # Hits (617190 total) | % Hits | Rank (847 total) | # Hits (437422 total) | % Hits | Rank (877 total) |
| CP4, CP7 | 7, 17 | Airway | Pseudomonas | 2399 | 3.2744 | 8 | 124 | 2.0622 | 5 | 938 | 0.1520 | 40 | 102178 | 23.3591 | 2 |
| CP8, CP13, CP17, CP20, CP26 | 20, 15, 17, 19, 17 | Airway | Bacillus | 1567 | 2.1388 | 10 | 6 | 0.0998 | 82 | 44156 | 7.1544 | 3 | 1145 | 0.2618 | 28 |
| CP9 | 24 | Airway | Neisseria | 329 | 0.4491 | 28 | 2 | 0.0333 | 142 | 44 | 0.0071 | 170 | 633 | 0.1447 | 41 |
| CP18, CP19 | 8, 15 | Airway | Brevibacillus | 0 | 0.0000 | - | 0 | 0.0000 | - | 23 | 0.0037 | 233 | 22 | 0.0050 | 243 |
| B3 | 6 | Liver | Saccharothrix | 0 | 0.0000 | - | 0 | 0.0000 | - | 225 | 0.0365 | 218 | 2 | 0.0005 | 699 |
| B8 | 2 | Liver | Halobacillus | 5 | 0.0068 | 263 | 0 | 0.0000 | - | 11 | 0.0018 | 344 | 5 | 0.0011 | 498 |
| B9 | 22 | Spleen | Lactobacillus | 232 | 0.3167 | 36 | 30 | 0.4989 | 28 | 6183 | 1.0018 | 16 | 5360 | 1.2254 | 9 |
| B11 | 23 | Liver | Aerococcus | 3 | 0.0041 | 333 | 0 | 0.0000 | - | 99 | 0.0160 | 115 | 4 | 0.0009 | 546 |
| B17 | 7 | Spleen | Klebsiella | 14 | 0.0191 | 181 | 5 | 0.0832 | 91 | 4 | 0.0006 | 539 | 281 | 0.0642 | 63 |
| B21 | 21 | Spleen | Lysobacter | 203 | 0.2771 | 40 | 1 | 0.0166 | 202 | 0 | 0.0000 | - | 3 | 0.0007 | 603 |
| B29 | 18 | Spleen | Streptomyces | 23 | 0.0314 | 138 | 2 | 0.0333 | 142 | 276 | 0.0447 | 73 | 117 | 0.0267 | 107 |
| B30 | 13 | Spleen | Planomicrobium | 4 | 0.0055 | 295 | 0 | 0.0000 | - | 1 | 0.0002 | 847 | 6 | 0.0014 | 458 |
| B38 | 2 | Spleen | Pediococcus | 2 | 0.0027 | 393 | 0 | 0.0000 | - | 10 | 0.0016 | 363 | 3 | 0.0007 | 603 |

**Table S2 Representation of CP genera in mouse airway microbiome profiles.** Genus-level phylogenetic assignments of the CPs, and of the B strains, were compared to those of mouse airway microbiome constituents previously identified in four independent 16S rRNA profiling efforts[21–25]. For each CP and B genus, listed are: 1) The number of times it is represented in a given airway microbiome profile ("# Hits"); 2) Its relative abundance in the profile ("% Hits"); and 3) Its relative rank within the profile, with the most abundant genus in the profile designated as "Rank" = 1. For each profile, the total number of 16S rRNA sequences placed at the genus level, and the total number of genera represented, are listed the headings of the "# Hits" and "Rank" columns, respectively.

| Isolate | Source | | Species | Profile 1 (Krogfelt) | | | Profile 2 (Klinman) | | | Profile 3 (Schöler) | | | Profile 4 (Dickson) | | |
|---|---|---|---|---|---|---|---|---|---|---|---|---|---|---|---|
| | Mouse | Tissue | | # Hits (60364 total) | % Hits | Rank (1358 total) | # Hits (5631 total) | % Hits | Rank (275 total) | # Hits (370596 total) | % Hits | Rank (5357 total) | # Hits (282127 total) | % Hits | Rank (4361 total) |
| CP4 | 7 | Airway | fluorescens | 6 | 0.0099 | 449 | 2 | 0.0355 | 179 | 23 | 0.0062 | 946 | 819 | 0.2903 | 32 |
| CP7 | 17 | Airway | stutzeri | 17 | 0.0282 | 237 | 1 | 0.0178 | 275 | 6 | 0.0016 | 2185 | 234 | 0.0829 | 92 |
| CP8 | 20 | Airway | velezensis | 2 | 0.0033 | 906 | 0 | 0.0000 | - | 0 | 0.0000 | - | 0 | 0.0000 | - |
| CP9 | 24 | Airway | lactamica | 0 | 0.0000 | - | 0 | 0.0000 | - | 1 | 0.0003 | 5357 | 0 | 0.0000 | - |
| CP13 | 15 | Airway | licheniformis | 12 | 0.0199 | 302 | 0 | 0.0000 | - | 26 | 0.0070 | 859 | 26 | 0.0092 | 521 |
| CP17 | 17 | Airway | megaterium | 88 | 0.1458 | 79 | 0 | 0.0000 | - | 7600 | 2.0508 | 10 | 9 | 0.0032 | 1133 |
| CP18 | 8 | Airway | frigoritolerans | 0 | 0.0000 | - | 0 | 0.0000 | - | 24 | 0.0065 | 908 | 7 | 0.0025 | 1369 |
| CP19 | 15 | Airway | borstelensis | 0 | 0.0000 | - | 0 | 0.0000 | - | 0 | 0.0000 | - | 1 | 0.0004 | 4361 |
| CP20 | 19 | Airway | clausii | 0 | 0.0000 | - | 0 | 0.0000 | - | 7 | 0.0019 | 1996 | 1 | 0.0004 | 4361 |
| CP26 | 17 | Airway | licheniformis | 12 | 0.0199 | 302 | 0 | 0.0000 | - | 26 | 0.0070 | 859 | 26 | 0.0092 | 521 |
| B3 | 6 | Liver | coeruleofusca | 0 | 0.0000 | - | 0 | 0.0000 | - | 0 | 0.0000 | - | 0 | 0.0000 | - |
| B8 | 2 | Liver | litoralis | 0 | 0.0000 | - | 0 | 0.0000 | - | 0 | 0.0000 | - | 1 | 0.0004 | 4361 |
| B9 | 22 | Spleen | amylolyticus | 0 | 0.0000 | - | 0 | 0.0000 | - | 0 | 0.0000 | - | 0 | 0.0000 | - |
| B11 | 23 | Liver | suis | 0 | 0.0000 | - | 0 | 0.0000 | - | 0 | 0.0000 | - | 0 | 0.0000 | - |
| B17 | 7 | Spleen | pasteurii | 0 | 0.0000 | - | 0 | 0.0000 | - | 0 | 0.0000 | - | 0 | 0.0000 | - |
| B21 | 21 | Spleen | niabensis | 0 | 0.0000 | - | 0 | 0.0000 | - | 0 | 0.0000 | - | 0 | 0.0000 | - |
| B29 | 18 | Spleen | collinus | 0 | 0.0000 | - | 0 | 0.0000 | - | 0 | 0.0000 | - | 0 | 0.0000 | - |
| B30 | 13 | Spleen | glaciei | 0 | 0.0000 | - | 0 | 0.0000 | - | 0 | 0.0000 | - | 0 | 0.0000 | - |
| B38 | 2 | Spleen | cellicola | 0 | 0.0000 | - | 0 | 0.0000 | - | 0 | 0.0000 | - | 0 | 0.0000 | - |

**Table S3 Representation of CP species in mouse airway microbiome profiles.** Species-level phylogenetic assignments of the CPs, and of the B strains, were compared to those of mouse airway microbiome constituents previously identified in four independent 16S rRNA profiling efforts[21–25]. For each CP and B species, listed are: 1) The number of times it is represented in a given airway microbiome profile ("# Hits"); 2) Its relative abundance in the profile ("% Hits"); and 3) Its relative rank within the profile, with the most abundant species in the profile designated as "Rank" = 1. For each profile, the total number of 16S rRNA sequences placed at the species level, and the total number of species represented, are listed the headings of the "# Hits" and "Rank" columns, respectively.

| CP Combination | Relative IC50 (95% CI) | Niche Index Fraction (NIF) | Figure |
| --- | --- | --- | --- |
| CP19/CP26 | $3.524x10^{-5}$ ($2.576x10^{-5}$ – $5.10x10^{-5}$) | 0.34 | Figure S4A |
| CP17/CP20 | 1501 (1071 – 1935) | 0.49 | Figure S4B |
| CP8/CP19 | $4.707x10^{-5}$ ($3.256x10^{-5}$ – $9.914x10^{-5}$) | 0.56 | Figure 4C |
| CP19/CP13 | 0.002126 (0.0007314 – 0.008040) | 0.72 | Figure 4D |
| CP8/CP13 | n.a. | 1.78 | Figure 4E |

**Table S4 Relative IC50 and Niche Index Fraction values for each CP combination tested.** Relative IC50 values are reported as IC50 (95% confidence interval). Non-inhibitory combinations have relative IC50 values listed as n.a. (not applicable).

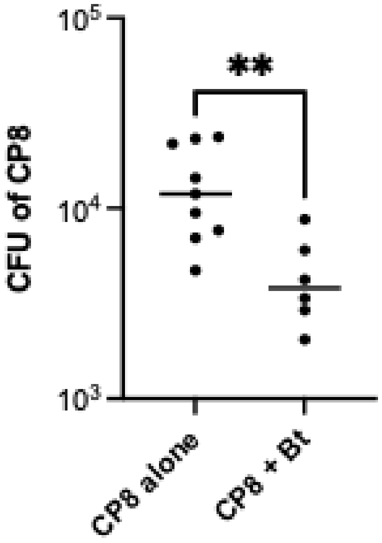


**Figure S6 CP8 colonization of the airway is significantly reduced in *Bt*-infected mice.** CP8 ($10^6$ CFU) was administered to the airway via OPA and after 3 days challenged with either PBS (CP8 alone) or *Bt* ($3x10^4$ - $5x10^5$ CFU) (CP8 + Bt) via OPA. Airway tissues were recovered at 3 days post-challenge (6 days post-CP8 administration) for enumeration of CP8 load (expressed as CFU per tissue homogenate). **P<0.002.

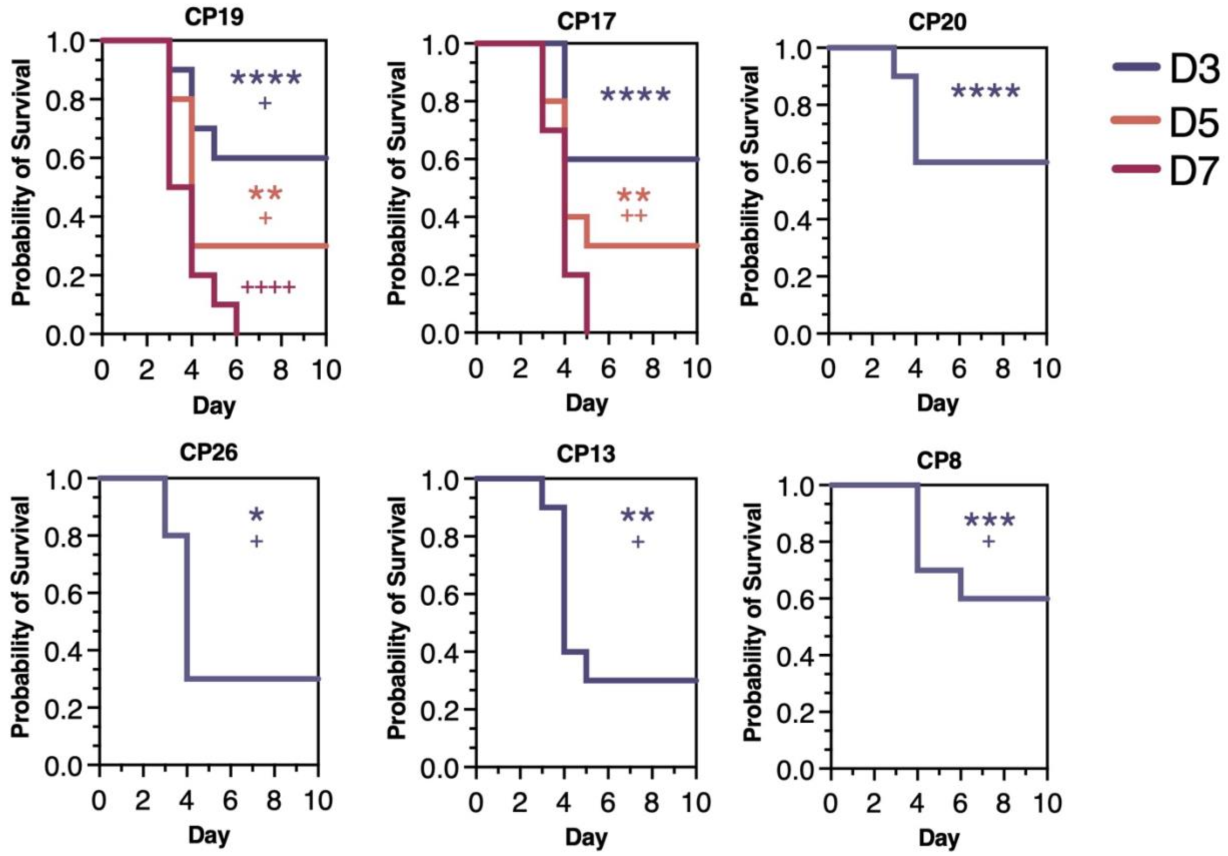


**Figure S7 Survival of mice treated with non-viable CPs prior to pathogen challenge.** CPs were rendered non-viable (via UV and heat treatment) and administered to the mouse airway (via OPA) at 3 days (blue), 5 days (orange), or 7 days (red) prior to challenge with *Bt* (via OPA). Survival was monitored for 10 days following pathogen challenge. For CP20, CP26, CP13, and CP8 treatment at 3 days prior to pathogen challenge was the only dosing regimen investigated because these CPs provided little to no protection when viable bacteria were administered at 5 days or 7 days prior to pathogen challenge (see Fig. 5E). Asterisks indicate significant differences between survival of mice treated with PBS (negative control) versus the non-viable CP, whereas plus signs indicate significant differences between survival of mice treated with the non-viable CP versus the viable CP, as calculated using the Holm-Šidák multiple comparisons test (****or $^{++++}P<0.0001$, ***or $^{+++}P<0.0002$, ** or $^{++}P<0.0021$, * or $^{+}P<0.0332$).

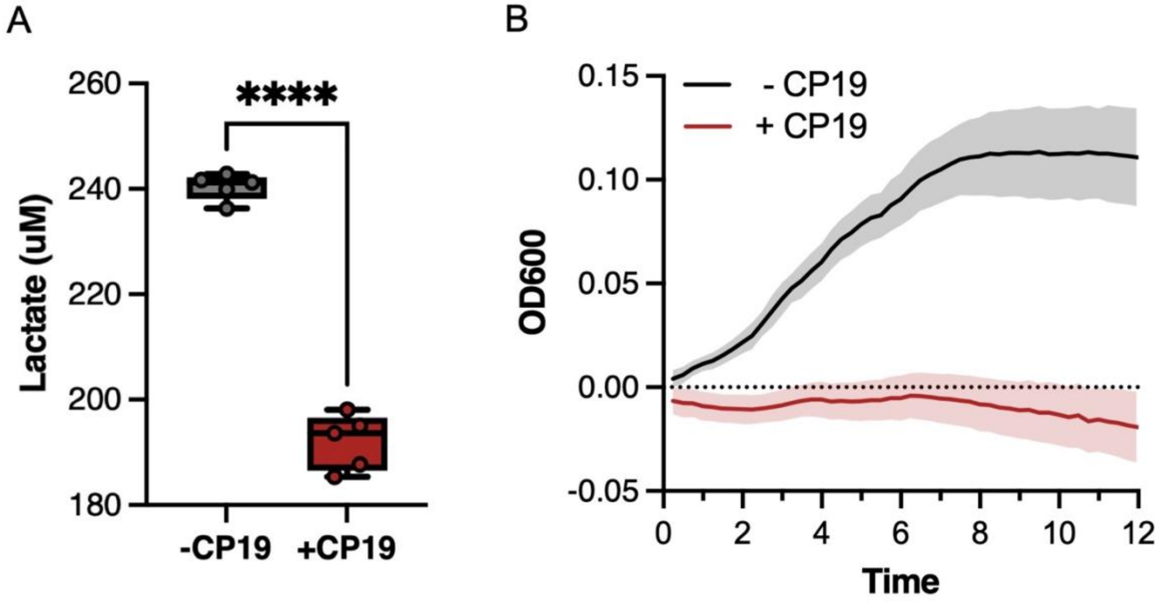


**Figure S8 CP19 consumption of lactate and inhibition of pathogen growth in mouse airway tissues *ex vivo*.** (A) Lactate levels in airway tissue homogenates treated with PBS (-CP19) or CP19 ($10^6$ CFU) (+CP19) for 24 hours. ****P<0.0001. (B) Growth of *Bt* in airway tissue homogenates pre-treated with PBS (-CP19) or CP19 ($10^6$ CFU) (+CP19) for 24 hours. Pre-treated tissue homogenates were filter sterilized, the filtrates were inoculated with *Bt* ($10^6$ CFU), and OD600 measurements were taken every 15 minutes to monitor growth of the pathogen over the course of 12 hours.
